# Using light-sheet microscopy to study spontaneous activity in the developing lateral-line system

**DOI:** 10.1101/2021.11.23.469686

**Authors:** Qiuxiang Zhang, Katie Kindt

## Abstract

Hair cells are the sensory receptors in the auditory and vestibular systems of all vertebrates, and in the lateral-line system of aquatic vertebrates. During development, spontaneous activity in hair cells shapes the formation of these sensory systems. In auditory hair cells of mice, coordinated waves of spontaneous activity can be triggered by concomitant activity in nearby supporting cells. But in mammals, developing auditory and vestibular hair cells can also autonomously generate spontaneous events independent of supporting cell activity. To date, significant progress has been made studying spontaneous activity in the auditory and vestibular systems of mammals, in isolated cultures. The purpose of this work is to explore the zebrafish lateral-line system as a model to study and understand spontaneous activity *in vivo*. Our work applies genetically encoded calcium indicators along with light-sheet fluorescence microscopy to visualize spontaneous calcium activity in the developing lateral-line system. Consistent with our previous work, we show that spontaneous calcium activity is present in developing lateral-line hair cells. We now show that supporting cells that surround hair cells, and cholinergic efferent terminals that directly contact hair cells are also spontaneously active. Using two-color functional imaging we demonstrate that spontaneous activity in hair cells does not correlate with activity in either supporting cells or cholinergic terminals. We find that during lateral-line development, hair cells autonomously generate spontaneous events. Using localized calcium indicators, we show that within hair cells, spontaneous calcium activity occurs in two distinct domains–the mechanosensory bundle and the presynapse. Further, spontaneous activity in the mechanosensory bundle ultimately drives spontaneous calcium influx at the presynapse. Comprehensively, our results indicate that in developing lateral-line hair cells, autonomously generated spontaneous activity originates with spontaneous mechanosensory events. Overall, with robust spontaneous activity three different cell types, the developing lateral line is a rich model to study these activities in an intact sensory organ. Future work studying this model may further our understanding of these activities and their role in sensory system formation, function and regeneration.

## Introduction

Spontaneous activity has been documented in developing sensory systems, including visual (Bansal et al., 2000;Warland et al., 2006;Ackman et al., 2012;Akrouh and Kerschensteiner, 2013), auditory (Tritsch et al., 2007;Johnson et al., 2011;Levic et al., 2011;Wang et al., 2015;Eckrich et al., 2018), vestibular (Holman et al., 2019) and somatosensory systems (Khazipov et al., 2004;Allene et al., 2008). Spontaneous activity in sensory cells can play a role locally in synapse maturation (Leighton and Lohmann, 2016) and ion channel expression (Moody and Bosma, 2005), as well as globally to pattern and refine downstream sensory circuits (Huberman et al., 2008;Clause et al., 2014;Newbold et al., 2020;Wosniack et al., 2021). For example, in the developing auditory system of mammals, spontaneous activity in sensory hair cells is thought to act locally help establish neuronal connections (Ceriani et al., 2019) and globally to shape the tonotopic maps along the auditory pathway (Babola et al., 2018). However, in mammals, the hair cells are enclosed in the bony structures of the ear–this location makes it difficult to access and record spontaneous activity *in vivo*. The purpose of this work is to explore the zebrafish lateral line as an *in vivo* model to study spontaneous activity in developing hair cells.

The most well-characterized studies of hair cell spontaneous activity have focused on the mouse auditory system (Kros et al., 1998;Marcotti et al., 2003;Wang et al., 2015;Ceriani et al., 2019;Babola et al., 2020;Jeng et al., 2020). During development, prior to the onset of hearing, spontaneous activity in auditory hair cells (inner and outer hair cells) takes two forms: 1) coordinated waves of activity among hair cells (Tritsch et al., 2007;Ceriani et al., 2019) and 2) uncoordinated, calcium action potentials generated autonomously (Kros et al., 1998;Marcotti et al., 2003;Johnson et al., 2011;Eckrich et al., 2018;Ceriani et al., 2019). For coordinated waves of activity among hair cells, concurrent waves of calcium propagate through glia-like supporting cells that surround or are adjacent to hair cells. In cochlear supporting cells, this activity is dependent on ATP and purinergic signaling and propagates via gap junction channels between supporting cells (Tritsch et al., 2007;Ceriani et al., 2019;Babola et al., 2021). Ultimately, ATP signaling in supporting cells drives spontaneous activity in nearby auditory hair cells. This coordinated activity is thought to be important for synapse refinement and for patterning the tonotopic axis along the cochlea and the downstream auditory pathway.

Independent of these coordinated waves of activity, both outer and inner auditory hair cells also fire spontaneous calcium action potentials during development (Kros et al., 1998;Marcotti et al., 2003;Johnson et al., 2011;Eckrich et al., 2018;Ceriani et al., 2019). These spontaneous action potentials are thought to be autonomously generated in hair cells (Johnson et al., 2012). Recent work has shown that this form of spontaneous activity may also be present in the vestibular system of mice (Holman et al., 2019) and in the lateral-line of zebrafish (Hiu-tung et al., 2019). In both mouse inner and outer auditory hair cells, and zebrafish lateral-line hair cells, Ca_V_1.3 calcium channels present at the hair cell presynapse are required for this form of spontaneous activity (Eckrich et al., 2018;Ceriani et al., 2019;Hiu-tung et al., 2019). Collectively, autonomously generated spontaneous activity appears to be a conserved feature of developing hair cell systems. Although the role of this activity is not fully understood, in the lateral line, we have shown that it can regulate presynapse size in developing hair cells (Hiu-tung et al., 2019). Autonomously generated spontaneous activity is also thought to be important for other aspects of hair cell and synapse maturation (Ceriani et al., 2019;Hiu-tung et al., 2019;Jeng et al., 2020). Currently the origin of autonomously generated spontaneous activity remains unclear.

In both the auditory and vestibular system of mice, in addition to supporting cells, cholinergic efferents physically interact with developing hair cells. These cholinergic efferents descend from the brainstem and synapse directly onto developing hair cells (Aschoff and Ostwald, 1987;Glowatzki and Fuchs, 2000;Roux et al., 2011). Furthermore, studies have shown that these efferents can modulate hair cell activity during development (Glowatzki and Fuchs, 2000;Katz et al., 2004;Holman et al., 2019). In addition, when the α9/α10 acetylcholine receptors required for this cholinergic modulation are disrupted, the patterns of hair cell spontaneous activity in the auditory epithelium are altered (Clause et al., 2014;Sendin et al., 2014). But whether these descending efferents are spontaneously active in hair cell systems and whether they can trigger spontaneous activity in developing hair cells is not known.

To study hair cells, supporting cells, and cholinergic efferents in the context of spontaneous activity, we explored the zebrafish lateral-line system. The lateral line is composed of superficial clusters of hair cells called neuromasts (Raible and Kruse, 2000). Unlike mammals, the lateral-line system can easily be accessed in intact zebrafish, making, it straightforward to visualize and study developing hair cells *in vivo*. With regard to development, the lateral line forms rapidly–when larvae are 2-3 days old (2-3 days post fertilization (dpf)), the majority of the hair cells are immature (Kindt et al., 2012). But when larvae are 5-6 days old (5-6 dpf), the majority of the hair cells are mature and the lateral-line system is functional (Kindt et al., 2012;Suli et al., 2012). This rapid developmental trajectory has made zebrafish an excellent system to study hair cell maturation. In addition to rapid development, zebrafish larvae are transparent, making it possible to visualize activity using genetically encoded calcium indicators (GECIs) (Lukasz and Kindt, 2018). The use of GECIs, along with advances in microscopy such as light-sheet fluorescence microcopy (LSFM) has increased the imaging potential of the zebrafish model (Vladimirov et al., 2014;Quirin et al., 2016). LSFM has been particularly advantageous–it uses plane illumination to rapidly image volumes with minimal photobleaching and phototoxicity even over extended periods of time (Power and Huisken, 2017).

Our study applies LSFM to study spontaneous calcium activity *in vivo,* in the zebrafish lateral-line system. We show that spontaneous calcium activities are present in several cell types in the periphery of the developing lateral line: hair cells, supporting cells, as well as cholinergic efferents that innervate hair cells. Using two-color functional imaging, along with pharmacology, we find that hair cell spontaneous activity occurs independent of activity in supporting cells and cholinergic efferents. Importantly, by using a membrane-localized calcium indicator in hair cells, we find that spontaneous calcium activity occurs in two distinct domains: the mechanosensory hair bundle and the presynaptic compartment. Further, our genetic and pharmacological analyses reveal that in lateral-line hair cells, spontaneous mechanosensory activity in the hair bundle drives spontaneous calcium influx at the presynapse. Thus, mechanosensory activity is the main source of spontaneous activity that is generated autonomously in hair cells. Overall, our study demonstrates that the lateral line is a valuable model to study many facets of spontaneous activity in an intact sensory system. The results of our study and future work using this *in vivo* system may improve our understanding of spontaneous activity and its role in sensory system formation and maintenance.

## Methods

### Zebrafish husbandry and strains

Zebrafish (Danio rerio) were raised at 28°C on a 14:10 h light/dark cycle. Larvae at 2-6 days post fertilization (dpf) were used for the experiments and were maintained in E3 embryo medium (in mM: 5 NaCl, 0.17 KCl, 0.33 CaCl_2_, and 0.33 MgSO_4_, buffered in HEPES pH 7.2) at a constant temperature of 28°C in an incubator. Because sex is not yet determined at these ages, we did not consider the animal’s gender in our research. Zebrafish work performed at the National Institute of Health was approved by the Animal Use Committee under animal study protocol #1362-13. Previously described mutant and transgenic zebrafish strains used in this study include: *Tg(myo6b:memGCaMP6s)^idc1^*(Jiang et al., 2017)*, Tg(myo6b:RGECO1)^vo10^* (Maeda et al., 2014)*, Tg(UAS:GCaMP6s)^mpn101^* and *Tg(Gal4:Chat)^mpn202^(Förster et al., 2017), pcdh15a^th263^(R306X)* (Seiler et al., 2005)*, and cav1.3a/cacna1d^tn004^ (R284C)* (Sidi et al., 2004).

### Vector construction and creation of transgenic lines

Plasmid construction was based on the tol2/gateway zebrafish kit (Kwan et al., 2007). To generate stable transgenic fish lines, plasmid DNA and tol2 transposase mRNA were injected into zebrafish embryos as previously described (Kwan et al., 2007). Using this approach, transgenic lines *Tg(she:GCaMP6s)^idc17^* and *Tg(myo6b:GCaMP6s)^idc18^* were created, using the supporting cell-specific promoter (*she*) (Quillien et al., 2017;Peloggia et al., 2021) or hair cell-specific promoter (*myo6b*) (Obholzer et al., 2008) to image calcium activity in the cytosol.

### Light sheet system construction and image acquisition

4D spontaneous calcium activity of zebrafish neuromast was imaged by a homebuilt dual-view inverted selective-plane illumination microscope (diSPIM). The microscope was built and aligned according to the previously described protocols (Kumar et al., 2014). Briefly, the bulk of the optomechanical hardware, automated stages, laser scanners, piezo elements for focus control, and control electronics were purchased from Applied Scientific Instruments (ASI, Eugene, Oregon, USA). Each arm of the diSPIM (SPIMA and SPIMB) consists of a water-dipping objective (Nikon CFI NIR Apo 40X Water DIC N2, Nikon, Melville, NY, USA) either emitting the laser light for excitation or collecting signals from the sample using an ORCA-Flash 4.0 sCMOS camera (Hamamatsu Photonics, Hamamatsu City, Shizuoka, Japan). Two optically pumped semiconductor laser (OBIS 488nm LX 30mW and OBIS 561nm LS 80mW, Coherent, CA, USA) were combined with a beam combiner (OBIS Galaxy, Coherent, CA, USA) and then routed to a fiber optic switch (eol 1×2 VIS, LEONI, VA, USA) for output laser wavelength selection. Two outputs from the fiber optic switch were fiber-coupled to the two diSPIM head laser scanners. The open-source platform Micro-Manager (https://micro-manager.org/) (Edelstein et al., 2010) was used for hardware interfacing, data capture, and storage. A TMC vibration isolation lab table (63P-9012M, TMC, Boston, MA) was used to house the light sheet system to minimize disturbance from external movement or vibration.

For single wavelength imaging, volumetric images were obtained at ∼0.33 Hz with 20 slices per volume (1 μm spacing) and 256 × 256 pixels per slice with 2 x2 binning to record the spontaneous calcium activity of zebrafish neuromasts over 15 mins. Two-color imaging was performed by sequentially imaging the green GECI and red GECI signals at ∼0.2 Hz with 8 slices per volume (2 μm spacing) over 15 mins. After fast volumetric calcium imaging, a high-resolution reference image stack was acquired from two views (SPIMA and SPIMB) respectively with 166 slices (0.5 μm spacing) and 1×1 binning. After high resolution imaging, images were cropped, and the backgrounds were subtracted in ImageJ (Schneider et al., 2012). Two image stacks from SPIMA and SPIMB views were then co-registered and deconvolved jointly to obtain a single volumetric image stack with isotropic spatial resolution and an isotropic voxel spacing (0.1625 μm/pixel x 0.1625 μm/pixel x 0.1625 μm/pixel). Custom software built in C++/CUDA(Guo et al., 2020) was used to conduct image registration and deconvolution on a graphics processing unit (GPU) card. The image registration process started with transforming the image stack from SPIMB with rotation, translation, or scaling and then overlaid and compared with those from SPIMA. By minimizing a cost function via Powell’s method (http://mathfaculty.fullerton.edu/mathews/n2003/PowellMethodMod.html), the best transformation matrix was obtained for registration. The images were then deconvolved jointly by an ‘unmatched back projector’ method, which could significantly accelerate deconvolution (Guo et al., 2020). The updated code published by Min Guo and Hari Shroff can be download from GitHub at: (https://github.com/eguomin/regDeconProject; https://github.com/eguomin/diSPIMFusion;https://github.com/eguomin/microImageLib).

The diSPIM resolution of our system was characterized using carboxylate-modified fluorescent beads (0.1 µm, F8803, Thermo Fisher Scientific, MA, USA) and diluted 100 times before use. The full width at half-maximum (FWHM) numbers were calculated for ∼8 beads along all axes before and after joint deconvolution (Kumar et al., 2014) (Supplementary Table 1). Before deconvolution, the FWHM of a bead was close to 0.5 μm (lateral) and 1.5 μm (axial). After registration of the two views (SPIMA and SPIMB) and joint deconvolution, the bead FWHM was approximately isotropic with 0.32 μm (lateral) and 0.41 μm (axial) resolution.

### Sample preparation for 4D spontaneous calcium imaging *in vivo*

Larvae at 2-6 dpf were first anesthetized with 0.03 % Ethyl 3-aminobenzoate methane sulfonate salt (Sigma-Aldrich, St. Louis, MO, USA) and pinned onto a Sylgard-filled recording chamber (I-2450, ASI, Eugene, Oregon, USA). After pinning, 125 μM α-Bungarotoxin (Tocris, Bristol, United Kingdom) was injected into the heart of intact larvae to suppress movement. Larvae were then rinsed with extracellular imaging solution (in mM: 140 NaCl, 2 KCl, 2 CaCl_2_, 1 MgCl_2,_ and 10 HEPES, pH 7.3) and allowed to recover. To image the spontaneous calcium activity, the larvae were kept in extracellular imaging solution without applying any external stimuli.

### Pharmacology

All drugs were prepared in extracellular solution with 0.1 % DMSO (except no DMSO was used with BAPTA). Animals were bathed in each drug (except BAPTA) for at least 15 mins prior to imaging. For BAPTA treatment, animals were incubated in BAPTA for 15 min, followed by a wash with extracellular solution. Listed below are the concentrations and incubation durations of the drugs used in this study:

| Drug | Mode of action | Vendor | Working<sup>207</sup><br>concentration |
| --- | --- | --- | --- |
| FFA | Gap junction blocker antagonist | Sigma-Aldrich | 25 μM |
| MRS2500 | P2yr1 selective antagonist | Tocris | 1 μM |
| Thapsigargin | Potent inhibitor of sarco/endoplasmic reticulum Ca<sup>2+</sup>-ATPases (SERCA) | Sigma-Aldrich | 250 nM |
| α-bungarotoxin (α-Btx) | Potent inhibitor of α9 or α10 nicotinic acetylcholine receptor (nAChR) | Tocris | 10 μM |
| Apamin | SK channel blocker | Tocris | 10 μM |
| Isradipine | L-type Ca<sup>2+</sup> channel CaV1.3a antagonist | Sigma-Aldrich | 10 μM |
| BAPTA-tetrasodium salt | Disrupts hair-bundle tip links | Thermofisher | 5 mM |

### Temporal quantification of 4D calcium imaging stacks

4D image stacks were first registered by the application of ImageJ macro, Correct 3D Drift (Arganda-Carreras et al., 2006;Parslow et al., 2014) and then converted into 2D time-series images by using the ImageJ Maximum Intensity Z-projection function. Another ImageJ macro, StackReg (Thevenaz et al., 1998) was used to further correct the movement artifact.

To quantify the average magnitude and frequency of spontaneous calcium activity, the maximum intensity 2D time-series images were processed in MATLAB R2020a (Mathworks, Natick, MA). First, a circular ROI was placed on each cell (diameter of ROI: ∼ 5 µm hair cell base; ∼ 2 µm hair bundle and ∼ 3 µm supporting cell soma). The fluorescent intensity value within each ROI was obtained for each time point. Then, the baseline of the temporal curves for each ROI was corrected by using the MATLAB imerode function. Next, the baseline corrected values were transformed into ΔF/F_0_, where the F_0_ was obtained for each time point by the MATLAB imerode function. Then, all values of ΔF/F_0_ less than 10 % were removed. Values below 10 % were considered noise; therefore, signals above 10 % ΔF/F_0_ were our threshold value for a true GCaMP6s signal. In hair cells a 10 % threshold was confirmed by imaging spontaneous GCaMP6s signals in the presence of isradipine where no signals were observed (***Figure 5***). The averaged magnitude of spontaneous activity per cell was obtained by dividing the integral or sum of GCaMP6s signals (ΔF/F_0_ >10 %) during the whole recording period by the total frames. Next, from our GCaMP6s recordings, peaks were detected by using MATLAB findpeaks function. The mean peak magnitude, frequency and duration were calculated per cell over the whole recording period (15 mins). Then the average magnitude, frequency, and duration per neuromast were computed by averaging measurements obtained from all the hair cells or supporting cells within the neuromast.

### Spatial visualization GCaMP6s signals in image stacks

To better observe the fluorescence intensity changes or the spontaneous calcium activity, fluorescence images were scaled and represented using color maps (***Figure 2***), with red indicating an increase in calcium signal relative to the resting period (baseline). The color maps were then superimposed into the grayscale morphological images for visualizing the spatial fluorescence intensity changes relative to the locations of specific hair cells or supporting cells within a neuromast.

The procedure to obtain these color maps has been described previously (Zhang et al., 2016). In ***Figure 2–figure supplement 2***, the spatial spontaneous activity images of the supporting cells were obtained by subtracting each image from baseline (reference image just prior to the rise signal) to represent the relative change in fluorescent signal from ΔF. In ***Figure 2***, the spatial spontaneous activity images of the supporting cells and hair cells were obtained by computing the averaged magnitude (ΔF/F_0_) over the whole recording period (15 min) pixel by pixel. The accompanying cytoGCaMP6s traces in ***Figure 2*** were obtained by placing an ROI over the entire neuromast during the pre- and post-drug incubation time windows.

### Statistical analysis

All data were analyzed and plotted with Prism 8 (Graphpad, San Diego, CA, USA). Values in the text and data with error bars on graphs and in text are expressed as mean ± SEM. All experiments were performed on a minimum of 3 animals, 4 neuromasts and on two independent days. These numbers were adequate to provide statistical power to avoid both Type I and Type II error. Datasets were confirmed for normality using a D’Agostino-Pearson omnibus normality test. Statistical significance between two conditions was determined by either an unpaired or paired t-test, or a Kruskal-Wallis test as appropriate.

## Results

### Fast, volumetric, *in vivo* imaging of spontaneous calcium activity in developing hair cells

Our previous work demonstrated that spontaneous presynaptic calcium activity in developing lateral-line hair cells is important for proper presynapse formation (Hiu-tung et al., 2019), yet how presynaptic calcium influx was initiated remained unclear. To study spontaneous activity during development in more detail, we constructed a dual-view inverted selective plane illumination microscope (diSPIM) (Kumar et al., 2014). This light-sheet microscope is specialized for fast, continuous imaging of volumes with isotropic resolution in XYZ, over long-time windows, with minimal photodamage or bleaching.

We tested the capability of this fluorescence microscope to detect spontaneous calcium activities in transgenic zebrafish lines expressing the cytosolic GECIs cytoGCaMP6s or cytoRGECO1 in hair cells. Using this approach, we observed robust spontaneous calcium signals in immature hair cells (day 3; ***Figure 1b1-b3, Figure 1d-d’, Movie 1, 2***). Consistent with our previous results where we used a membrane-localized indicator, memGCaMP6s to monitor spontaneous presynaptic calcium activity, cytoGCaMP6s signals were dramatically reduced when hair cells matured (day 6; ***Figure 1d-d’***). We found that this microscope system was suitably fast for imaging the time course of these calcium signals in volumes (83 µm x 83 µm x 20 µm volumes acquired within 100 ms every 3 s) that encompass the entire neuromast organ. Despite continuous, volumetric imaging over long-time windows (15 mins), we observed minimal-to-no photodamage or bleaching (***Figure 1b3***). Overall, our data indicates that the diSPIM microscope is well-suited to rapidly measure spontaneous calcium activities within entire neuromast organs.

**Figure 1:**
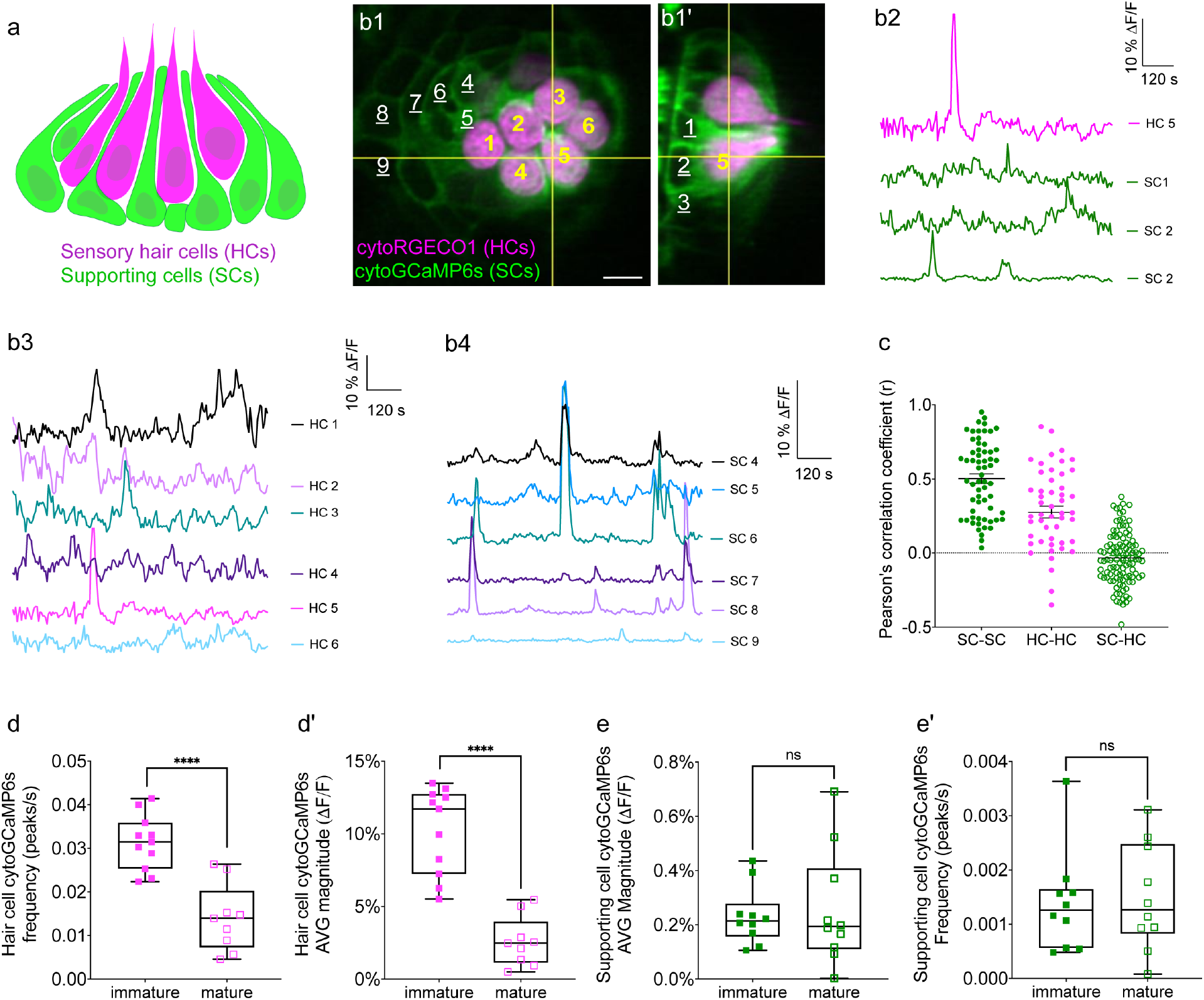
Simultaneous imaging of spontaneous activities in hair cells and supporting cells. (a) Cartoon of a neuromast organ illustrating sensory hair cells (magenta) and the surrounding supporting cells (green). (b1-b1’) Images of a representative, day 3, double transgenic neuromast expressing cytoRGECO1 (hair cells, numbered in yellow) and cytoGCaMP6s (supporting cells, a subset is numbered in white, underscored). A top-down view (b1) and the corresponding side view of its orthogonal projection (b1’) clearly show the hair cells and the surrounding supporting cells at day 3. (b2) Temporal curves of spontaneous calcium activity in hair cell 5 (labeled in b1-b1’) and its 3 surrounding supporting cells (labeled in b1’) within the 15 min recording window show distinct time courses. (b3) Temporal curves of spontaneous calcium activity in the cluster of 6 hair cells (labeled in b1) reveals distinct response profiles. (b4) Temporal curves of spontaneous calcium activity in the 7 supporting cells (labeled in b1) reveals some correlation and synchronization among neighboring supporting cells. (c) Comparisons of Pearson’s R of spontaneous calcium activities at day 3 in neighboring supporting cells, neighboring hair cells and hair cells with its surrounding hair cells, n = 6 neuromasts for HC-HC or SC-SC R values and n = 4 neuromasts for HC-SC R values. (d-d’) Box plots showing quantification of the average magnitude (d) and frequency (d’) of spontaneous calcium activity in immature (day 3) and mature (day 6) hair cells. (e-e’) Box plots showing quantification of the average magnitude (e) and frequency (e’) of spontaneous calcium activity in supporting cells in mature (day 6) and immature (day 3) neuromasts. Each dot in d-e’ represents one neuromast. An unpaired t-test was used in (d-e’). **** p<0.0001. Scale bar = 5 μm.

**Figure 2.**
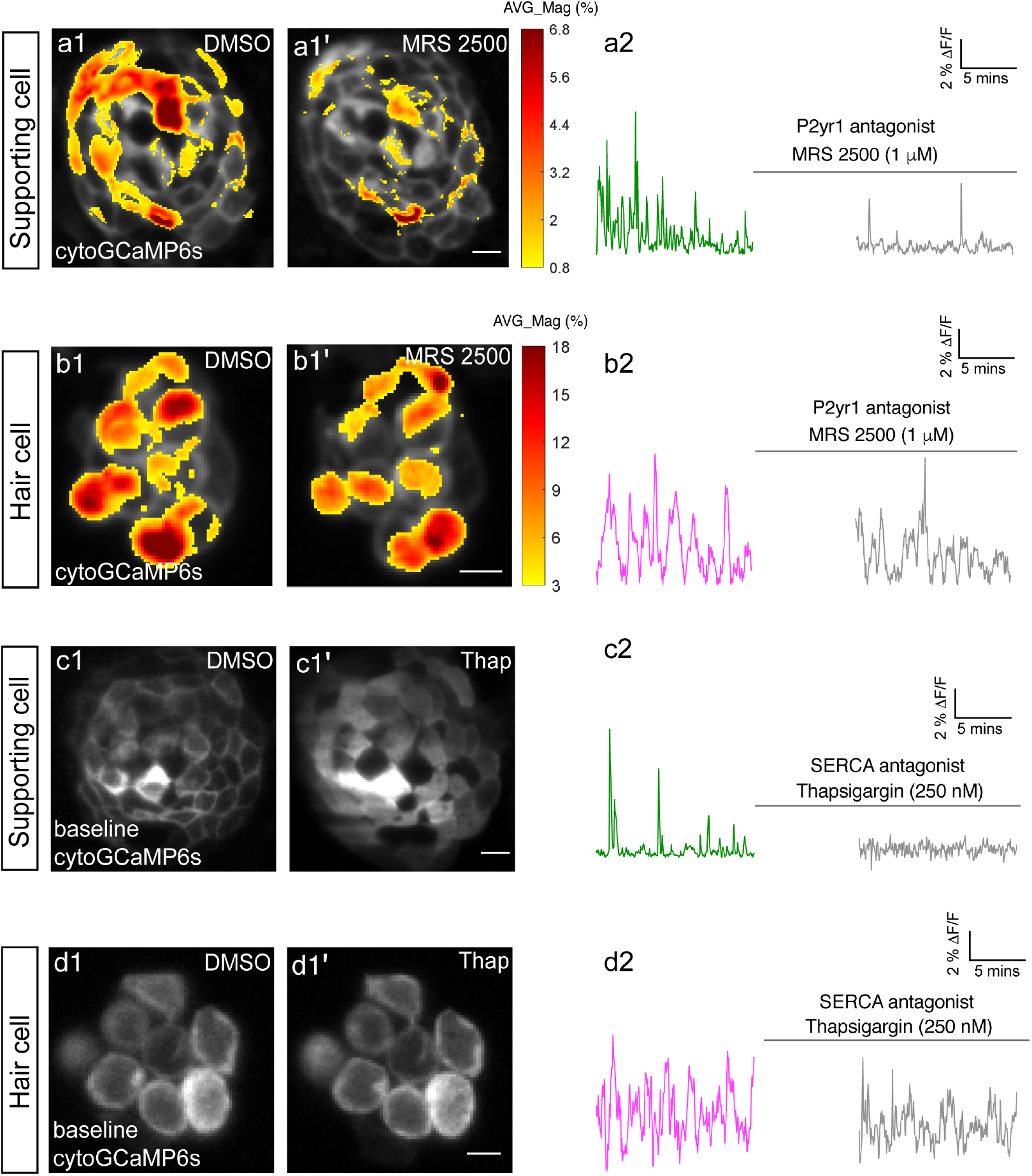
P2yr1 signaling is required for spontaneous activity in supporting cells but not in hair cells. The spatial patterns of the mean spontaneous calcium activities of the supporting cells (a1-a1’) or hair cells (b1-b1’) in DMSO and after 15 mins of treatment with 1 µM MRS2500. Measurements were performed in immature neuromasts at day 3. The ΔF/F GCaMP6s signals were averaged over each 900 s interval (pre- and post-treatment) and then colorized according to the heat map and superimposed onto a baseline image. The corresponding temporal curves of the mean signal magnitude across the whole neuromast in supporting cells (a2) and in hair cells (b2) in DMSO and after 15 mins of treatment with 1 µM MRS2500. The cytosolic baseline calcium in the supporting cells (c1-c1’) and hair cells (d1-d1’) in DMSO and after 15 mins of treatment with 250 nM Thapsigargin. Measurements were performed in immature neuromasts at day 3. The corresponding temporal curves of the mean signal magnitude across the whole neuromast in supporting cells (c2) and in hair cells (d2) in DMSO and after 15 mins of treatment with 250 nM Thapsigargin. Scale bar = 5 μm.

**Figure 3.**
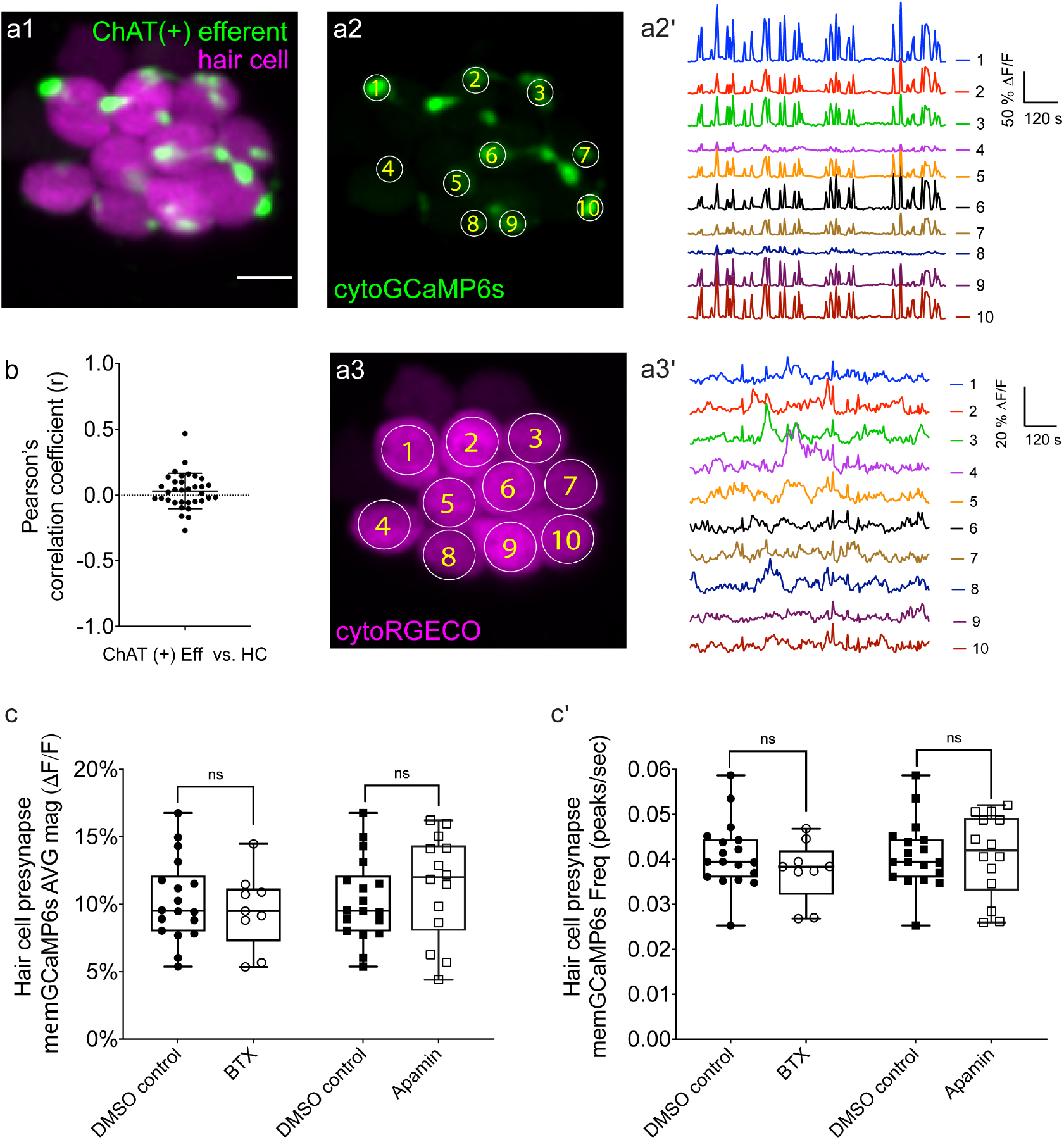
Two-color imaging of spontaneous activities in hair cells and cholinergic efferent terminals. (a1) A representative maximum intensity z-projection image of a double-transgenic zebrafish line expressing the red GECI cytoRGECO1 in hair cells and the green GECI cytoGCaMP6s in the cholinergic efferents terminals at day 3. (a2) Image depicting cholinergic efferent terminals with individual terminals contacting different hair cells in a1 are labeled accordingly. (a2’) The corresponding temporal curves of spontaneous calcium activities of the 10 efferent terminals indicated in a2. (a3) Image of the individual hair cells in a1 labeled according to innervating efferent terminal. (a3’) The corresponding temporal curves of spontaneous calcium activities of the 10 hair cells indicated in a3. (b) Pearson’s R values of spontaneous calcium activity between each hair cell and its contacting cholinergic efferent terminal, n = 34 hair cell-efferent terminal pairs. Box plots showing the average magnitude (c) and frequency (c’) of spontaneous calcium activity at the hair cell presynapse in DMSO and after the treatment with 10 μM α-Btx and 10 μM apamin. Each point in c-c’ represents one neuromast. All measurements were performed in immature neuromasts at day 3. A paired t-test was used in c-c’. Scale bar = 5 μm.

### Spontaneous calcium activity is present in zebrafish supporting cells

In cultured explants of the mammalian inner ear work has demonstrated that, in addition to hair cells, the glia-like supporting cells that surround auditory and vestibular hair cells also display spontaneous calcium activity (Wang et al., 2015;Eckrich et al., 2018;Ceriani et al., 2019;Holman et al., 2019). Furthermore, in the developing auditory system, waves of spontaneous activity in supporting cells can trigger spontaneous activity in hair cells (Wang et al., 2015;Ceriani et al., 2019;Babola et al., 2020;Babola et al., 2021). Like mammalian hair cell systems, hair cells in the zebrafish lateral line are isolated and surrounded by supporting cells (***Figure 1a-b’***). Therefore, we tested whether zebrafish supporting cells were spontaneously active during development.

To measure spontaneous calcium activity, we created a transgenic zebrafish line expressing the cytosolic GECI cytoGCaMP6s in supporting cells (***Figure 1b1-b1’***). Using this line, we observed that in developing neuromasts, spontaneous calcium signals were also present in supporting cells (day 3; ***Figure 1b2, b4***; ***Figure 1e-e’*,*Movie 3***). We compared the properties of the spontaneous signals detected in supporting cells with those detected in the hair cells. Our analyses revealed that the average magnitude of cytoGCaMP6s signal above baseline during the 15-min recording window (average magnitude; ***Figure 1—figure supplement 1a***) was much higher in hair cells than that in supporting cells. Furthermore, there were also significantly more spontaneous calcium events occurring in the hair cells compared with supporting cells (peaks per second; ***Figure 1—figure supplement 1a’***). In addition to more spontaneous calcium events, each spontaneous calcium event lasted much longer in the hair cells compared to supporting cells (duration; ***Figure 1—figure supplement 1a’’***). Despite more events with a longer time course, on average, the peak magnitudes were comparable between hair cells and supporting cells (average peak mean; ***Figure 1—figure supplement 1a’’’***). These data indicate that in the zebrafish lateral-line system, spontaneous calcium activity is present in both hair cells and supporting cells.

In the mammalian auditory and vestibular systems, spontaneous activity in supporting cells is thought to be restricted to development (Wang et al., 2015;Ceriani et al., 2019;Holman et al., 2019). Interestingly, we found no significant difference in spontaneous cytoGCaMP6s signals in supporting cells when we compared immature (day 3) to mature (day 6) neuromast organs (***Figure 1e-e’***). Overall, our work indicates that spontaneous activity is present in both hair cells and supporting cells in the zebrafish lateral line. Interestingly, although hair cell spontaneous activity occurs primarily during development (***Figure 1d-d’***), in supporting cells, this activity is retained upon sensory system maturation.

### Spontaneous calcium activities in hair cells and supporting cells do not coincide

To understand if, similar to the mammalian auditory system, spontaneous calcium activities in hair cells and supporting cells are linked, we performed two-color (red and green) calcium imaging. Here we simultaneously imaged the activity in both hair cells and supporting cells, using a double-transgenic zebrafish line expressing the red cytosolic GECI cytoRGECO1 in hair cells (Sheets et al., 2017) along with the green cytosolic GECI cytoGCaMP6s in supporting cells (***Figure 1b1-b1’***). We focused our analyses on immature neuromasts (day 3) when spontaneous calcium activity is robust in hair cells. Using our diSPIM microscope, along with these calcium reporters, we were able to simultaneously detect spontaneous calcium activity in all hair cells and supporting cells within whole neuromast organs (***Figure 1b-c, Figure 1—figure supplement 2, Movie 4***).

Next, we investigated whether there was a relationship between the spontaneous calcium activities in hair cells and the surrounding supporting cells during development (day 3). In addition to the rapid volumetric calcium imaging, diSPIM light-sheet systems can acquire 3D image stacks that can be viewed in a top-down view (***Figure 1b1***) or a corresponding side view (***Figure 1b1’***) with comparable resolution (Kumar et al., 2014). This isotropic 3D resolution enabled us to spatially delineate all hair cells, as well as all surrounding supporting cells (***Figure 1b1-b1’, Figure 1—figure supplement 2a1-a1’, b1-b1’, c1-c1’***). Using this approach, we observed that the spontaneous calcium activities in hair cells and neighboring supporting cells did not coincide. For example, the side-view image in ***Figure 1b1’*** clearly highlights a hair cell (HC 5) and its three surrounding supporting cells (SC 1-3). By plotting the temporal curves of these 4 cells (***Figure 1b2***), we found that both hair cells and the supporting cells were spontaneously active. But the temporal response profile in the hair cell was distinct from its surrounding supporting cells (***Figure 1—figure supplement 2c1-c3***; Pearson’s R = 0.19, 0.19 and -0.01). Similar results were obtained for other hair cell and supporting-cell pairings in this example (***Figure 1—figure supplement 2a1-b3***). We extended this analysis and calculated the Pearson’s correlation coefficients from cell pairings across many developing neuromasts organs (day 3) and overall found no correlation in spontaneous calcium activities between hair cells and surrounding supporting cells (***Figure 1c***, Pearson’s R = -0.04 ± 0.02, n = 4 neuromasts). In summary, analysis of our two-color functional imaging in hair cells and supporting cells revealed that there was little to no correlation in spontaneous calcium activities between immature hair cells and surrounding supporting cells.

After determining that there was no correlation in spontaneous calcium activities between hair cells and supporting cells, we examined whether there was a correlation among populations of hair cells or supporting cells within neuromast organs. Within a given neuromast, we observed that each hair cell showed distinct response profiles (example, ***Figure 1b3***) and the correlation between any two neighboring hair cells was low (example, ***Figure 1—figure supplement 2d1-d2***). When we examined spontaneous calcium activity between neighboring hair cells across many developing neuromast organs, quantification revealed little to no correlation (***Figure 1c***, Pearson’s R = 0. 28 ± 0.04, n = 6 neuromasts). In contrast to hair cells, by plotting temporal curves of spontaneous calcium activity from individual supporting cells, we observed clear examples of synchronized activities (examples, ***Figure 1b4, Figure 1—figure supplement 2d1, d3*)**. When examining the spontaneous calcium activity between neighboring supporting cells, quantification revealed a relatively high correlation (***Figure 1c***, Pearson’s R = 0.50 ± 0.03, n = 6 neuromasts). Numerous studies have demonstrated that supporting cells in mammals (Mulroy et al., 1993;Lautermann et al., 1998;Sato et al., 1998;Kikuchi et al., 2000) and zebrafish (Zhang et al., 2018) are electrically coupled via gap junction channels. Therefore, one possible reason for synchronized activities between supporting cells are these gap junction channels. Consistent with this idea, we found that blockage of gap junctions with flufenamic acid (FFA) significantly reduced the correlation in spontaneous calcium activity between neighboring supporting cells (control: Pearson’s R = 0.45 ± 0.03; 10 µM FFA: Pearson’s R = 0.08 ± 0.02, n = 4 neuromasts).

Together, our results suggest that there is no obvious temporal relationship between spontaneous calcium activities in hair cells and supporting cells. Furthermore, distinct temporal properties indicate the source or mechanism underlying these two distinct spontaneous calcium signals may also be different.

### P2yr1-ER signaling underlies spontaneous calcium activity in supporting cells but not in hair cells

In the mammalian auditory system, extracellular ATP and P2 purinergic signaling is required for spontaneous activity in supporting cells (Babola et al., 2020;Babola et al., 2021). Previous work has shown that pharmacological block of P2RY1 receptors and downstream phospholipase C-β (PLC-β) signaling components blocked spontaneous activity in supporting cells (Babola et al., 2020). Therefore, we used pharmacology to investigate whether P2RY1 signaling is required for initiating spontaneous calcium activity in supporting cells within the zebrafish lateral-line system.

For our analyses we measured cytoGCaMP6s signals in supporting cells in developing neuromasts. Using this approach, we found that the application of P2RY1 receptor antagonist MRS2500 significantly reduced the magnitude and frequency of spontaneous cytoGCaMP6s signals in zebrafish supporting cells (example, day3, ***Figure 2a1-a2***; ***Figure 2—figure supplement 1a-a’***). This result indicates that, like the mammalian auditory system, P2yr1 is required for spontaneous activity in zebrafish supporting cells. Numerous studies have shown that P2RY1 signaling activates phospholipase C-β (PLC-β), leading to the cleavage of phosphatidylinositol 4,5-bisphosphate (PIP2) into diacylglycerol and inositol-1,4,5-trisphosphate (IP_3_) (Bucheimer and Linden, 2004). IP_3_ binds to IP_3_ receptors on the ER, resulting in the release of calcium from the ER into the cytoplasm (Desai and Leitinger, 2014). This conserved signaling cascade was shown to be present in supporting cells in the developing auditory epithelium (Babola et al., 2020). To determine if ER calcium was released downstream of P2ry1 in zebrafish supporting cells, we examined spontaneous calcium activity before and after inhibitors of ER calcium release. For our analysis, we used the Sarco/ER calcium ATPase (SERCA) blocker, thapsigargin. SERCA block prevents reentry of calcium to the ER from the cytosol and ultimately depletes ER calcium stores (Courjaret et al., 2018). After application of thapsigargin, we observed that the baseline cytoGCaMP6s levels increased dramatically in supporting cells within 5 minutes (example, day3, ***Figure 2c1-c1’***). In addition to changes in baseline calcium, we also observed that the application of thapsigargin dramatically reduced spontaneous calcium activity in supporting cells (example, day3, ***Figure 2c2***).

To further support the idea that ER release from the cytosol acts downstream of P2yr1 receptors, we examined the spatiotemporal properties of spontaneous calcium activity in supporting cells more closely. For our analysis, we imaged supporting cells in cross-section longitudinally. We acquired cytoGCaMP6s signals in a single plane, at fast image acquisition speed (10 fr/s compared to 1 volume/3 s in ***Figure 1*** and ***Figure 2***). We used heat maps to illustrate the spatial increase in calcium signals during a representative spontaneous event (***Figure 2—figure supplement 2a1***). We also plotted the calcium signals at 3 distinct positions within the cell (***Figure 2—figure supplement 2a2-a4:*** *top, middle, and bottom).* Both the heat maps and plots revealed that spontaneous calcium activity in the supporting cells initiates in the upper regions of the cells. These plots revealed that within 0.5 s after initiation, a calcium increase was observed throughout the entire supporting cell, with activity at the top of the cell detected ∼300 ms before activity at the bottom of the cell. This apical initiation is interesting because it is the proposed location of the purinergic receptors present in supporting cells within the mammalian auditory system (Babola et al., 2021).

Lastly, we also tested whether P2yr1 or SERCA block altered spontaneous calcium activity in developing hair cells. Using cytoGCaMP6s, we found that P2yr1 block with MRS2500 did not impact spontaneous calcium activity in immature hair cells (example, day 3, ***Figure 2b1-b2, Figure 2—figure supplement 1b-b’***). Similarly, we examined the impact of SERCA block on spontaneous calcium activity in immature hair cells. We found that neither the baseline calcium (***Figure 2d1-d1’***) nor the spontaneous calcium signals (***Figure 2d2***) were significantly altered after thapsigargin application. These data indicate that neither P2yr1 receptor function nor ER calcium stores are critical for spontaneous calcium activity in immature hair cells.

Overall, we found that similar to the mouse auditory system, in the zebrafish lateral line, a P2yr1-ER signaling cascade is required for spontaneous calcium activity in supporting cells. While the pharmacological block of P2yr1 or ER calcium stores dramatically blocks spontaneous calcium activity in supporting cells, this same block does not alter spontaneous calcium activity in zebrafish hair cells. These pharmacological results indicate that during development, zebrafish hair cells and supporting cells use distinct mechanisms to generate spontaneous calcium signals. In addition, these results provide further evidence that in the zebrafish lateral line, spontaneous calcium activity in hair cells does not require concomitant activity in surrounding supporting cells.

### Two-color imaging reveals no link between spontaneous activities in hair cells and efferents

If supporting cells do not trigger spontaneous calcium activity in hair cells in zebrafish, then how is this activity generated in developing hair cells? In both the mouse auditory and vestibular system, cholinergic efferent fibers descend from the brainstem and synapse onto developing hair cells (Aschoff and Ostwald, 1987;Glowatzki and Fuchs, 2000;Roux et al., 2011;Johnson et al., 2013). These efferent contacts have been shown to regulate the activity of developing hair cells (Glowatzki and Fuchs, 2000;Katz et al., 2004;Holman et al., 2019). In the developing zebrafish lateral line, cholinergic efferents also synapse directly onto immature hair cells (Zhang et al., 2018;Freixas et al., 2021) (***Figure 3a1***). But whether these efferents are spontaneously active or whether they regulate the spontaneous calcium activity in immature zebrafish lateral-line hair cells is not known.

To measure calcium signals in cholinergic efferents, we used a transgenic line expressing green cytoGCaMP6s *UAS:cytoGCaMP6s* driven by *Gal4:Chat* (Förster et al., 2017). We used this transgenic line in combination with a transgenic line expressing red cytoRGECO1 in hair cells (***Figure 3a1***) to monitor calcium activities in these two cell types simultaneously (***Movie 5***). Using this approach, we found that in developing neuromast organs, without any external stimuli, the efferent terminals contacting hair cells were spontaneously active (examples, day3, ***Figure 3a1-a2’, figure 3—figure supplement 1a1-a3’***). In addition, we observed that among all efferent terminals within a neuromast, spontaneous calcium activities were highly synchronized (***Figure 3—figure supplement 1b,*** Pearson’s R = 0.92 ± 0.007, n = 4 neuromasts). While the magnitude and frequency of spontaneous calcium activity in efferents terminals was robust during development (day 3), upon lateral-line maturation (day 6) the activity was reduced (***Figure 3—figure supplement 1c-c’***). Because spontaneous calcium activities in both efferent terminals and hair cells were largely restricted to development, we reasoned that these activities could be linked. But when we compared the spontaneous calcium activity in each efferent terminal with its hair cell target, we found no correlation (***Figure 3b***; Pearson’s R = 0.03 ± 0.02, n = 34 hair cell-efferent terminal pairs). This lack of correlation indicates that cholinergic efferent neurons may not directly trigger spontaneous calcium activity in hair cells.

To further examine the role of cholinergic efferents in hair cell spontaneous activity, we took a pharmacological approach. Cholinergic efferents share a molecular mechanism that is conserved among vertebrate hair cell systems (Glowatzki and Fuchs, 2000;Hiel et al., 2000;Parks et al., 2017;Carpaneto Freixas et al., 2021). Acetylcholine released from these efferents acts on α9 or α10 nicotinic acetylcholine receptors (nAChR), which allows calcium to enter near the hair cell presynapse. This calcium influx activates calcium-dependent potassium SK channels leading to hyperpolarization of the hair cell. To determine if this cascade of events impacts hair cell spontaneous activity, we applied α-bungarotoxin (α-Btx) and apamin, compounds that block α9 nAChRs and SK channels respectively. Recent work has shown that these compounds are effective at disrupting efferent neurotransmission onto lateral-line hair cells (Carpaneto Freixas et al., 2021). Because cholinergic signaling is thought to impact the hair cell presynapse, for our analyses we used a membrane-localized indicator memGCaMP6s which we have used previously to specifically measure spontaneous presynaptic calcium activity (Hiu-tung et al., 2019). We found that neither α-Btx nor apamin impacted spontaneous presynaptic calcium activity in immature hair cells (day 3, ***Figure 3c-c’***). Overall, our imaging revealed that both hair cells and efferents have spontaneous calcium activities during development. Further, our 2-color imaging and pharmacological results support the conclusion that cholinergic efferents are not required for spontaneous calcium activity in hair cells.

### Spontaneous calcium activity occurs in the mechanosensory bundle and at the presynapse

Our two-color calcium imaging (***Figure 1a-c, Figure 3a1-a3’***) indicates that in the zebrafish lateral line, neither supporting cells nor cholinergic efferent neurons–two cell types that are spontaneously active and directly contact hair cells–trigger hair cell spontaneous calcium activity. Therefore, we hypothesized that spontaneous calcium activity in hair cells may be generated autonomously. In mature hair cells the main pathway leading to calcium influx are generated in response to sensory stimuli. In response to stimuli, mechanotransduction (MET) channels open, leading to a cationic influx in the mechanosensory hair bundle that can trigger opening of presynaptic calcium channels at the presynapse (Pickles et al., 1984). (***Figure 4a***). Therefore, one possibility is that spontaneous MET channel activity also triggers opening of presynaptic calcium channels during development. While, we previously demonstrated that spontaneous calcium activity is present at the presynapse of immature lateral-line hair cells (Hiu-tung et al., 2019), whether there is spontaneous calcium activity in mechanosensory hair bundles was not known.

**Figure 4.**
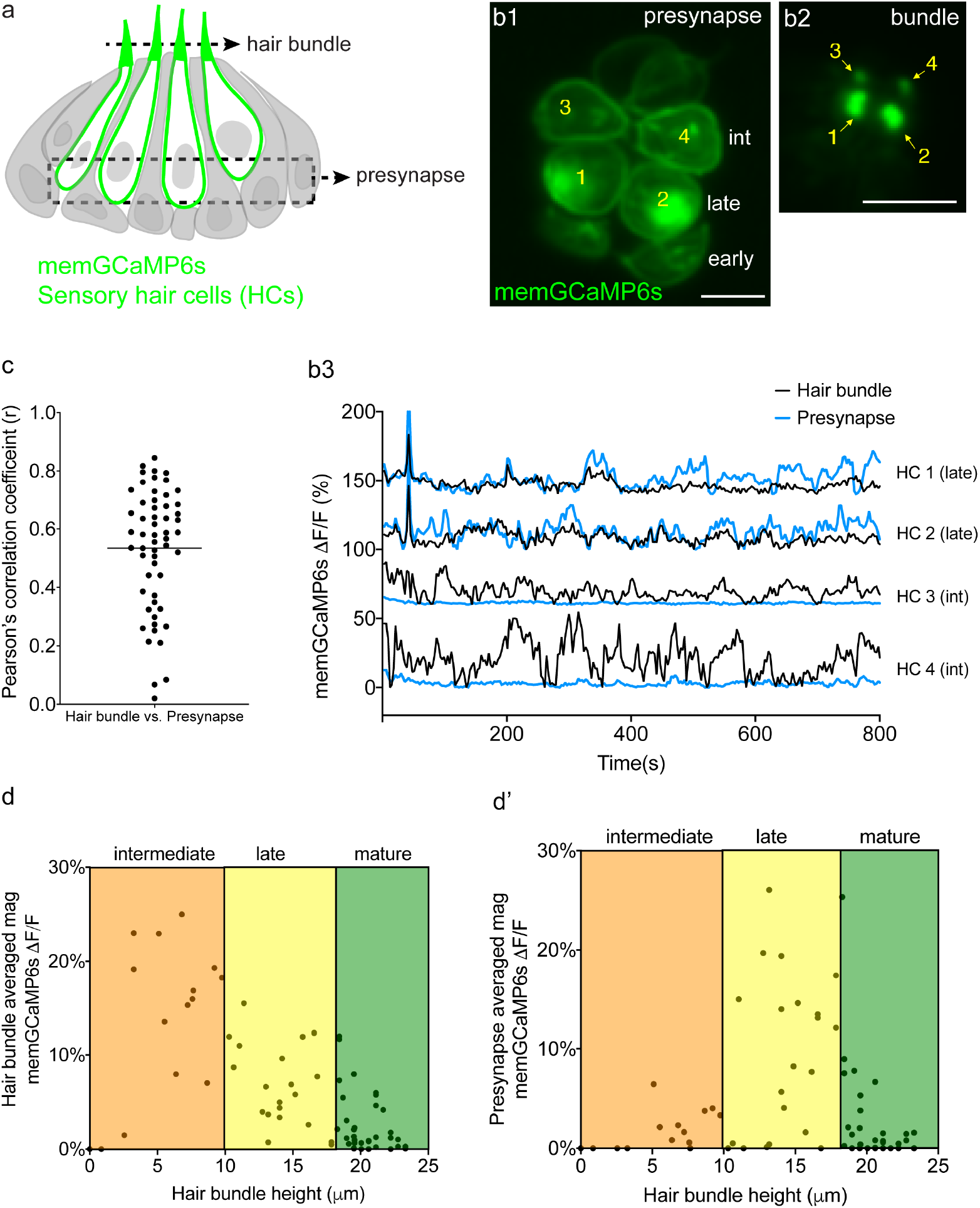
Spontaneous calcium activity occurs in the mechanosensory bundle and at the presynapse. (a) Cartoon of a neuromast organ illustrating hair cells (green) expressing memGCaMP6s, surrounded by supporting cells (gray). (b1) Image of a representative neuromast showing the hair cell presynaptic region indicated by the dashed box in ‘a’ taken from immature hair cells at day 2. Hair cells at 3 different developmental stages are labeled: early (no detectable hair bundle), intermediate (3,4), and late (1,2). (b2) An image of the hair bundles from the same cells as b1 through the plane indicated by the dash line in ‘a’. (b3) Paired temporal curves of spontaneous calcium activity in hair bundles (b2 black) and the presynaptic regions (b1 blue) taken from the same hair cells. (c) Pearson’s R values of spontaneous calcium activity between hair bundles and the presynaptic regions of the same sets of immature hair cells, n = 56 hair cells, day 2 and 3. (d-e) The average magnitude of spontaneous calcium activity in hair bundles and presynaptic region in hair cells during development. Hair bundle height is used to delineate the progressive stages: intermediate, late and mature. (d) The spontaneous calcium activity in hair bundles peaks early in development and decreases upon maturation (increasing hair bundle height). (d’) The spontaneous calcium activity in the presynaptic regions peaks after activity in the hair bundle and decreases upon maturation. Scale bar = 5 μm.

To measure spontaneous calcium activity in mechanosensory bundles, we used a membrane-localized GECI memGCaMP6s (Jiang et al., 2017). Previously we have used memGCaMP6s to measure evoked calcium signals in the mechanosensory bundle as well as both evoked and spontaneous calcium influx at the presynapse (Zhang et al., 2018;Hiu-tung et al., 2019). Using memGCaMP6s, in the absence of mechanical stimuli, we observed robust, spontaneous calcium activity in apical mechanosensory bundles (***Movie 6***). We examined spontaneous calcium activity in mechanosensory bundles in immature (day 3) and mature (day 6) hair cells. We found that similar to measurements using cytoGCaMP6s (***Figure 1d-d’***) and memGCaMP6s examining presynaptic activity (Hiu-tung et al., 2019) (***Figure 4—figure supplement 1b-b’***), spontaneous activity in mechanosensory bundles was largely restricted to immature hair cells (***Figure 4—figure supplement 1a-a’***).

By selecting top-down views of our 3D images, we were able to isolate and simultaneously measure activities in both a basal, presynaptic plane (example, ***Figure 4b1-b3***) and an apical mechanosensory bundle plane (example, ***Figure 4b2***) within the same hair cells. When we examined individual hair cells, in many cases, we found that the spontaneous calcium activity in each apical mechanosensory bundle and its respective presynaptic region were highly correlated (example, ***Figure 4b3***, see HC 1 and HC 2). In other cases, we observed hair cells with spontaneous calcium activity in the mechanosensory bundle but not in the presynaptic region (example, ***Figure 4b3***, see HC 3 and HC 4). On average, the correlation in spontaneous calcium activity at the apex and the base of individual hair cells was high in immature hair cells (***Figure 4c***, day 2, Pearson’s R = 0.60 ± 0.035, n = 32 hair cells; day 3 Pearson’s R = 0.54 ± 0.027, n = 56 hair cells). This relatively high correlation indicates that spontaneous calcium activity at the presynaptic region may be driven by or be related to spontaneous calcium activity in the mechanosensory bundle.

We looked more closely at individual hair cells to understand why we observed a correlation in spontaneous activity in the presynaptic compartment and mechanosensory bundle in some hair cells but not in others. What we found is that the strength of the correlation depends on the developmental stage of the hair cells. Within a developing neuromast (day 2-3), there are hair cells at different developmental stages (Kindt et al., 2012;Dow et al., 2018). Previous work has demonstrated that in the lateral line, hair cell stage can be estimated by measuring the height of the tallest structure in the mechanosensory bundle, the kinocilium (Stage-hair bundle height: early- not detectable; intermediate- 1-10 µm; late- >10<18 µm; mature- >18 µm. Lateral-line hair cells take roughly 20 hrs to complete this maturation) (Kindt et al., 2012;Dow et al., 2018). Overall, spontaneous calcium activity in both the mechanosensory bundle and the presynaptic compartment increased in strength during development and was dramatically reduced upon hair cell maturation (***Figure 4d-d’***). In addition, we observed that, on average, spontaneous calcium activity in the mechanosensory bundle initiated at an earlier development stage compared to activity at the presynapse (***Figure 4d-d’***). This differential time course resulted in a lack of correlation between these two signals in hair cells during the earliest stages of development. For example, in younger hair cells, we observed spontaneous calcium activity in apical mechanosensory bundles but not in the basal presynaptic compartment (labeled as ‘int’ in ***Figure 4b1***; see HC 3 and HC 4 in ***Figure 4b1-b3***). In slightly more mature hair cells, the spontaneous calcium activity in the hair bundle and presynapse were highly correlated (labeled as ‘late’ in ***Figure 4b1***; see HC 1 and HC2 in ***Figure 4b1-b3***).

Overall, by measuring calcium activities at the membrane, we demonstrate for the first time that spontaneous calcium activity is present in both apical mechanosensory bundles and basal presynaptic compartments *in vivo*. At the early stages of hair cell development, spontaneous calcium activity is present in mechanosensory bundles but not at the presynapse. At later stages of hair cell development, spontaneous calcium activities in mechanosensory bundles and the presynaptic compartments are highly correlated.

### Ca_V_1.3 channels are required for spontaneous presynaptic- but not hair bundle-activity

Our memGCaMP6s-based calcium imaging indicates that spontaneous activity in immature hair cells occurs in two distinct subcellular compartments. But how these signals are initiated in lateral-line hair cells remained unclear. Work in mammals and zebrafish indicates that hair cells can initiate spontaneous calcium activity autonomously during development. In the mouse auditory system and in the zebrafish lateral line, these potentials rely on Ca_V_1.3, a L-type calcium channel present at the hair cell presynapse (Eckrich et al., 2018;Ceriani et al., 2019). Therefore, we used both genetics and pharmacology to examine whether Ca_V_1.3 channels are required for spontaneous calcium activity in hair cells of the lateral line.

For our analyses, we examined activity in *ca_V_1.3a* zebrafish mutants and in wildtype animals after treatment with isradipine, a Ca_V_1.3 channel antagonist. Both of these manipulations have been shown to block evoked presynaptic calcium influx in the hair cells of mammals and zebrafish (Sheets et al., 2012;Huang and Moser, 2018;Zhang et al., 2018). To measure spontaneous calcium activity, we used memGCaMP6s to detect signals in the mechanosensory bundle and at the presynapse. We found that in *ca_V_1.3a* mutants or after acute block of Ca_V_1.3 channels with isradipine, spontaneous calcium activity in the mechanosensory bundle was not significantly changed compared to controls (***Figure 5a-d, Figure 5—figure supplement 1a-b’***). However, we found that spontaneous calcium activity occurring at the hair cell presynapse was abolished in *ca_v_1.3a* mutants (***Figure 5e-g, Figure 5—figure supplement 1c-d’***) and after isradipine treatment (***Figure 5h, Figure 5—figure supplement 1c-d’***). Together these manipulations indicate that spontaneous calcium activity can occur in the mechanosensory bundle without accompanying activity at the presynapse. Furthermore, they demonstrate that while Ca_v_1.3 channels are not required for spontaneous calcium activity in mechanosensory bundles, they are essential for spontaneous calcium activity at the presynapse.

**Figure 5.**
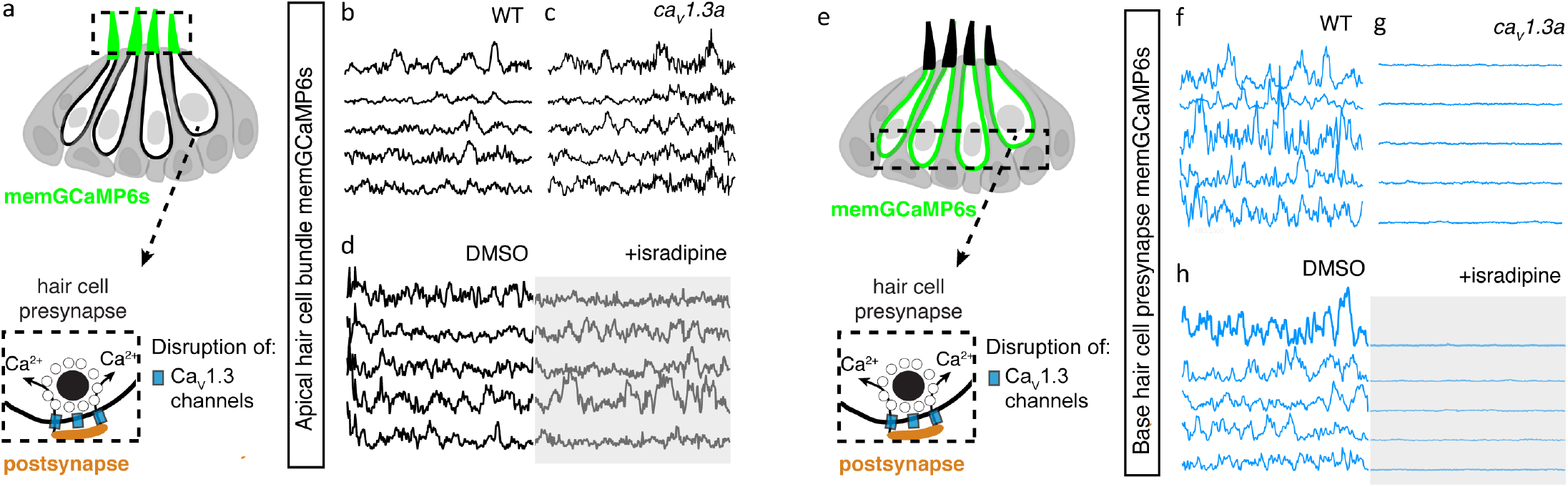
Spontaneous calcium activity at the presynapse requires Ca_V_1.3 channels. (a) Diagram illustrating the hair cells with the hair bundles highlighted by a dashed box where we measure their spontaneous calcium activity. The structure of a single ribbon synapse at the base of a hair cell is illustrated, as well as the calcium channels beneath it. (b-d) Representative memGCaMP6s temporal traces show that disruption of Ca_V_1.3a channels using *cav1.3a* mutants (b-c) or pharmacological block using 10 µM isradipine (d) does not impact spontaneous activity in apical hair bundles. (e) Diagram illustrating the hair cells with the presynaptic area highlighted by a dashed box where we measure the spontaneous presynaptic calcium activity. (f-h) Representative memGCaMP6s temporal traces show that disruption of calcium channels using *cav1.3a* mutants (f-g) or pharmacological block using 10 µM isradipine (h) completely blocked spontaneous activity at the hair cell presynapse. All measurements were performed in immature hair cells at day 3.

### Mechanotransduction in hair bundles is required for spontaneous activity at the presynapse

Our functional calcium imaging indicates that spontaneous calcium activity at the presynapse, when present, occurs concomitantly with activity in the mechanosensory bundle (***Figure 4c***). Additionally, in mature hair cells, evoked activity opens MET channels leading to an influx of cations, including calcium, that triggers opening of Ca_V_1.3 channels at the presynapse (Bechstedt and Howard, 2007). Therefore, we hypothesized that the spontaneous opening of MET channels in mechanosensory bundles might trigger activity at the presynapse. To test this hypothesis, we used genetics and pharmacology to determine whether MET channel function is required to trigger spontaneous calcium activity at the presynapse of immature hair cells.

For our genetic analysis, we used the zebrafish mutants lacking Protocadherin 15a (PCDH15), a core component of hair cell tip-link, a structure required to gate MET channels in mature hair cells (Seiler et al., 2005;Müller, 2008) (***Figure 6a***). For our pharmacological analysis, we applied BAPTA, an extracellular calcium chelator that breaks tip links and acutely disrupts MET channel function (Assad et al., 1991;Goodyear and Richardson, 2003). We found that MET channel disruption in *pcdh15a* mutants or after BAPTA treatment significantly reduced spontaneous calcium activity in mechanosensory bundles (***Figure 6a-d, Figure 6—figure supplement 1a-b’***). This indicates that in the lateral line, both evoked calcium activity in mature mechanosensory bundles and spontaneous calcium activity in developing mechanosensory bundles depends on MET function. Importantly, we found MET channel disruption also dramatically blocked spontaneous calcium activity at the hair cell presynapse (***Figure 6e-h, Figure 6—figure supplement 1c-d’***). After our manipulations we did observe some residual spontaneous calcium activity in the mechanosensory bundle and at the presynapse, which may reflect incomplete block of MET channel function. Overall, this result indicates that spontaneous activity at the presynapse is primarily driven by the spontaneous opening of MET channels. Thus, it is spontaneous MET that generates spontaneous calcium activity at the presynapse, in order to shape synapse formation and propagate information downstream to the developing sensory system.

**Figure 6.**
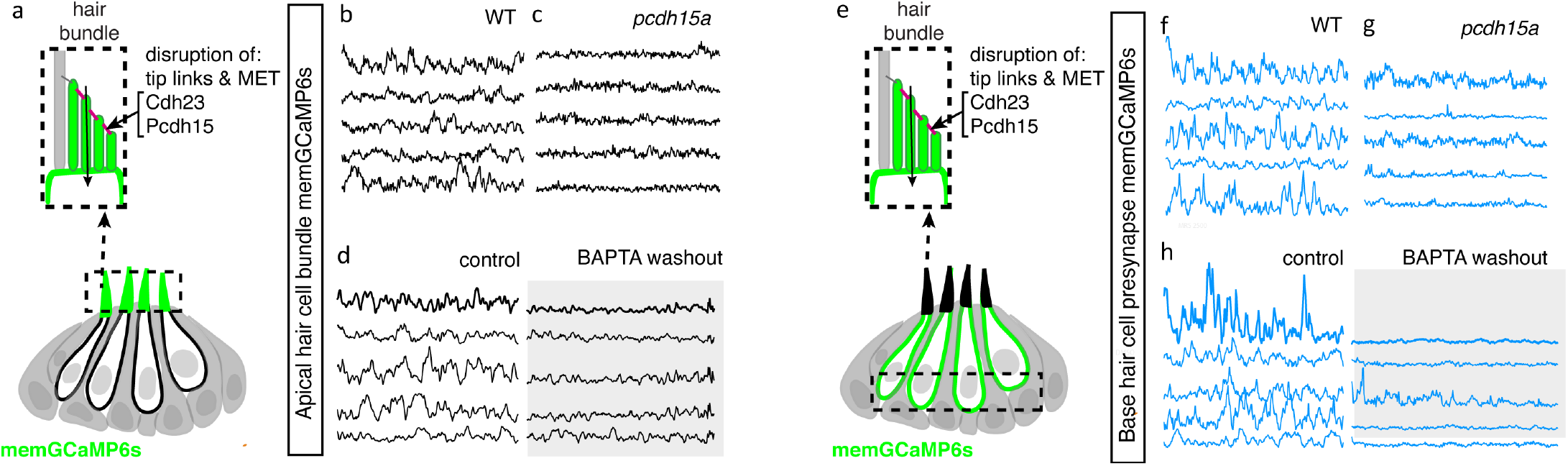
Mechanotransduction in hair bundles is required for spontaneous activity at the presynapse. (a) Diagram illustrating the hair cells with the hair bundles highlighted by a dashed box where we measure their spontaneous calcium activity. The structure of the hair bundle from a single hair cell at is illustrated on top. The mechanotransduction (MET) channels are gated by tip links made up of Pcdh15a Cdh23. Pharmacological application of BAPTA, an extracellular calcium chelator, breaks tip links and acutely disrupts MET function. (b-d) Representative GCaMP6s temporal traces show that disruption of tip links using *pcdh15a* mutants (b-c) or by pharmacologically disrupting tip links using BAPTA (d) disrupts spontaneous activity in hair bundles. (e) Diagram illustrating the hair cells with the hair cell presynaptic area highlighted by a dashed box at the base of the hair cells where we measure the spontaneous presynaptic calcium activity. (e-h) Representative GCaMP6s temporal traces show that disruption of tip links using *pcdh15a* mutants (f-g) or by pharmacologically disrupting tip links using BAPTA (h) also disrupts spontaneous calcium activity at the presynapse. All measurements were performed in immature hair cells at day 3.

## Discussion

In this work, we show that zebrafish can be used as an *in vivo* model to study spontaneous activity in developing hair cell sensory systems. The origin and role of spontaneous activity in hair cell systems is an important area of research in sensory neuroscience. Our work uses genetics, pharmacology, GECIs along with cutting-edge LSFM to image and characterize spontaneous calcium activity in the lateral-line system. We find that spontaneous activity is present in hair cells, supporting cells and efferent terminals*–*spontaneous activity in each cell type occurs with largely distinct developmental, temporal and mechanistic origins (***Figure 7***). Overall, our work demonstrates that the lateral line presents a robust model to study many facets of spontaneous activity, all within an intact sensory system.

**Figure 7.**
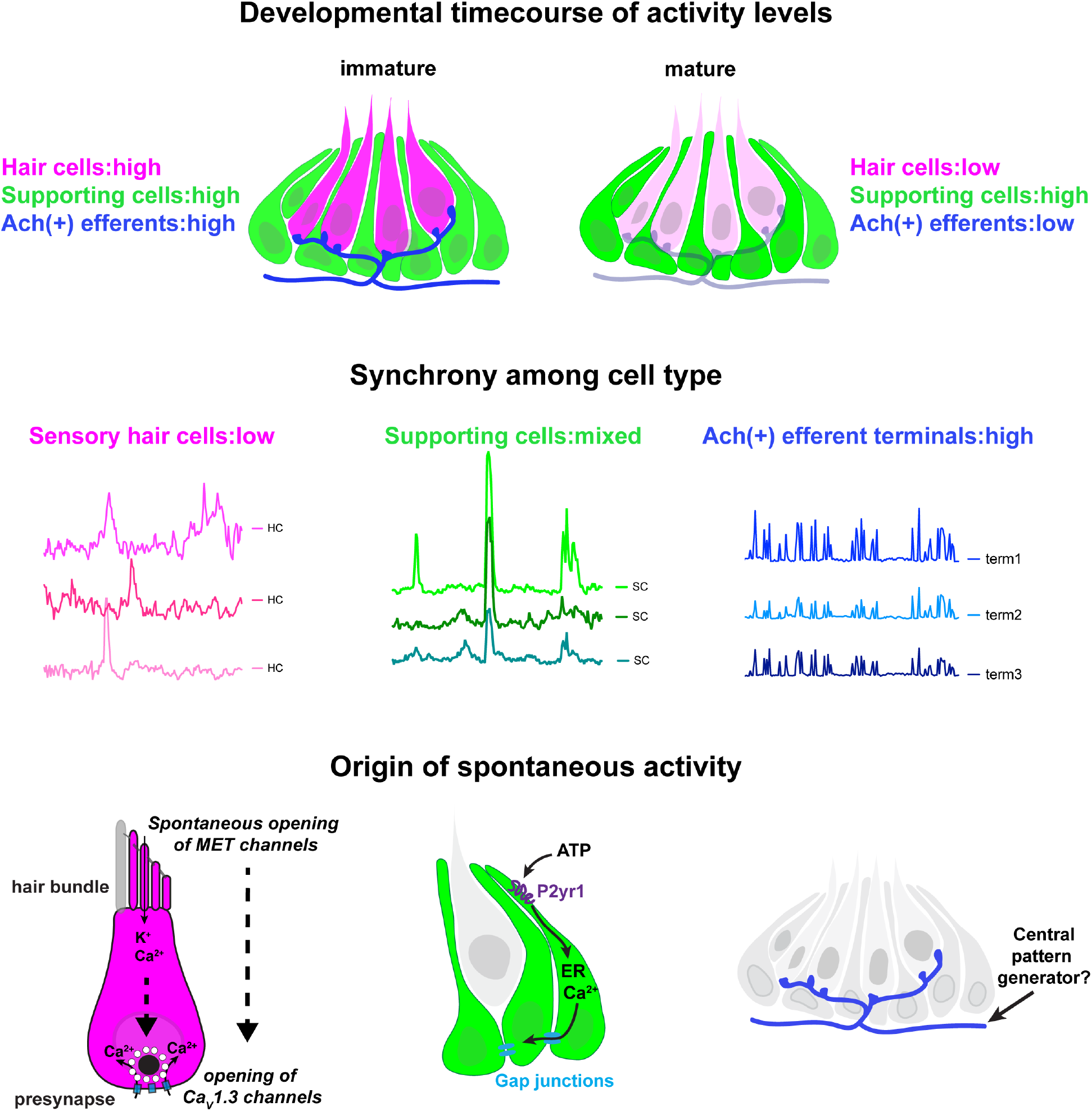
Locations, mechanisms and developmental timecourses of spontaneous calcium activities in the lateral-line system. Top panel: we reliably detected spontaneous calcium activity in three cell types within the lateral-line system: hair cells, supporting cells and cholinergic (Ach (+)) efferent neurons. Spontaneous calcium activity in the supporting cells and cholinergic efferents is not correlated with calcium signals in hair cells. During development spontaneous calcium signals are robust in all three cell types (left side). Upon maturation, spontaneous calcium signals are decreased in hair cells and efferent neurons, but maintained in supporting cells (right side) Middle panel: Within a neuromast spontaneous calcium signals among populations of hair cells are not correlated (left side); supporting cells are moderately correlated (middle); cholinergic terminals are highly correlated (right side). Bottom panel: In developing hair cells, spontaneous opening of MET channels leads to calcium signals in mechanosensory hair bundles. This apical activity triggers opening of Ca_V_1.3 channels at the presynapse, resulting in spontaneous presynaptic calcium signals (left panel). In supporting cells, extracellular ATP acts on P2yr1 receptors. P2yr1 signaling leads to calcium release from the ER, giving rise to a spontaneous calcium signal. Spontaneous calcium signals can be propagated to neighboring supporting cells via gap junction channels (middle panel). Spontaneous activity in cholinergic efferents may be coupled to activity in spinal motor neurons that is present during development in order to form the central pattern generator required for locomotion (right panel).

### The origin of spontaneous calcium activity in the zebrafish hair cells

In the developing mouse auditory system, spontaneous activity in supporting cells is linked to waves of activity in hair cells (Tritsch et al., 2007;Wang et al., 2015;Babola et al., 2020). In addition to supporting cells, cholinergic efferents are thought to shape hair cell activity in the developing mouse auditory system (Glowatzki and Fuchs, 2000;Katz et al., 2004;Johnson et al., 2013). In our study on the zebrafish lateral line, we find that spontaneous calcium signals in hair cells are still intact when supporting cell spontaneous activity is blocked (Block P2ry1 receptors or ER calcium release, ***Figure 2, Figure 2—figure supplement 1***) or when the cholinergic receptors on hair cells are blocked (Block α9 nAChRs, ***Figure 3c-c’***). This indicates, that in the zebrafish lateral line, neither supporting cells nor efferent neurons have a striking impact on spontaneous calcium activity in developing hair cells.

In both the developing mouse auditory and vestibular systems, hair cells also autonomously generate spontaneous calcium activity (Marcotti et al., 2003;Levic et al., 2011;Eckrich et al., 2018;Holman et al., 2019). Based on our work we hypothesize that the spontaneous calcium signals we observe in lateral-line hair cells are analogous to these autonomously generated signals. In mice these calcium signals have been visualized using either cytosolic GECIs or calcium dyes, or by recording calcium action potentials using electrophysiology (Marcotti et al., 2003;Levic et al., 2011;Eckrich et al., 2018;Holman et al., 2019). Our work uses both a cytosolic GECI (cytoGCaMP6s) as well as a membrane-localized GECI (memGCaMP6s) to visualize spontaneous calcium signals in lateral-line hair cells (***Figure 1*** and ***Figure 4***). Importantly, using memGCaMP6s, we find that spontaneous activity within hair cells occurs in two spatially distinct domains–the mechanosensory bundle and the presynapse (***Figure 4***). Using memGCaMP6s in lateral-line hair cells, we find that Ca_V_1.3 channels, present at the developing presynapse, are essential for spontaneous presynaptic calcium signals (***Figure 5e-h, Figure 5—figure supplement 1c-d’***). Related work in mouse auditory hair cells, has also found that Ca_V_1.3 calcium channels are important for spontaneous calcium signals that are autonomously generated (Brandt et al., 2003;Eckrich et al., 2018;Ceriani et al., 2019). Furthermore, both work in mice and zebrafish indicates that this Ca_V_1.3-dependent, presynaptic calcium activity is critical for proper hair cell synapse formation (Ceriani et al., 2019;Hiu-tung et al., 2019).

Interesting, our work found that disrupting Ca_V_1.3 channel function did not alter spontaneous calcium activity in mechanosensory bundles. This result supports the idea that spontaneous calcium activity in mechanosensory bundles occurs upstream of activity at the presynapse. In support of this idea, we found that 1) activity in these two domains is highly correlated (***Figure 4c-b3***) 2) at the earliest stages of hair cell development spontaneous activity was present in mechanosensory bundles but not at the presynapse (***Figure 4b3, d-d’***) and 3) blocking MET channel function significantly reduced spontaneous calcium activity at the presynapse (***Figure 6e-h, Figure 6—figure supplement 1c-d’***). Together these results demonstrate that in developing lateral-line hair cells, spontaneous MET activity in mechanosensory bundles likely triggers presynaptic calcium influx. Previous work in zebrafish has also shown the MET activity is major player that drives tonic or spontaneous release of vesicles at mature hair cell synapses (Trapani and Nicolson, 2011). Further, work in mice has shown that MET activity in developing hair cells is important for hair cell survival, hair cell synapse maintenance, and for the proper maturation of hair cells, afferent neurons and efferent neurons (Marcotti et al., 2006;Corns et al., 2018;Sun et al., 2018). Based on our work, and other studies, during development, it is clear that MET function in the mechanosensory bundle has many important roles. Our work shows that during development, spontaneous MET function is responsible for initiating spontaneous activity that is autonomously generated in lateral-line hair cells. This apical spontaneous activity ultimately leads to the opening of Ca_V_1.3 channels at the presynapse, likely triggering activity in the afferent neurons and downstream auditory pathway.

### The origin and role of spontaneous calcium activity in the zebrafish supporting cells

Similar to mouse auditory and vestibular systems, we detected robust spontaneous calcium activity in developing supporting cells within the lateral-line system (***Figure 1b1-b1’, b4***). In the mouse auditory system, spontaneous activity between neighboring supporting cells is coupled and occurs in waves; these waves trigger activity in nearby hair cells (Tritsch et al., 2007). These waves coordinate spontaneous activity among populations of hair cells to set up tonotopic (frequency) maps along the auditory pathway (Clause et al., 2014). In contrast to the auditory system, in the developing mouse vestibular system spontaneous activity in supporting cells does not occur in waves (Holman et al., 2019). In addition, no coupling is seen between neighboring supporting cells, and activity between hair cells and supporting cells is not coordinated. Like the mouse vestibular system, in the lateral line, we were unable to detect waves of spontaneous activity across neuromast organs. It is possible that there are no waves of activity in the vestibular and lateral-line systems because there is no need to coordinate activity among neighboring hair cells to set up tonotopic maps. Interestingly, unlike the mouse vestibular system, in the lateral line, we observed that activity between neighboring supporting cells was coupled (***Figure 1b4-c, Figure 2—figure supplement 2d1, d3***). The level of coupling was higher in the outer most layer of supporting cells (mantle cells) compared to the more central supporting cells (***Figure 2—figure supplement 3a-b***). In the future it will be interesting to examine spontaneous activity in different subsets of supporting cells within the mammalian vestibular system to see if coupling is limited to specific cell types, or to a particular region of the epithelium.

Together, our work in zebrafish along with work in mammals demonstrates that spontaneous calcium signals in supporting cells are a conserved feature among hair cell systems. In addition, our findings indicate that the molecular pathway required to trigger spontaneous calcium signals in supporting cells may also be partly conserved. Our work in zebrafish indicates a P2yr1 signaling cascade leads to calcium release from the ER; this calcium release gives rise to the spontaneous calcium signals in lateral-line supporting cells (***Figure 2a1-a2, c1-c2***). This is consistent with recent work in the mouse auditory epithelium that found autocrine P2yr1 signaling, followed by ER calcium release, is required for the spontaneous calcium activity observed in supporting cells (Babola et al., 2020;Babola et al., 2021). Further, studies have shown that application of ATP evokes calcium signals in supporting cells within the auditory and vestibular epithelium–these signals also require ER calcium release (Dulon et al., 1993;Holman et al., 2019;Rabbitt and Holman, 2021). Together these studies point towards a conserved P2-ER calcium axis that drives calcium signals in the supporting cells of hair cell epithelia. In the future, by using GECIs along with ATP sensors (Lobas et al., 2019) and two-color imaging, the zebrafish lateral line presents a useful model to study the dynamics of calcium and ATP *in vivo* during spontaneous events.

Although the molecular cascade giving rise to calcium signals in supporting cells appears largely conserved among hair cell systems, we did observe one striking difference in the lateral-line system. While spontaneous activity in supporting cells is transient in the mouse auditory and vestibular systems (Tritsch et al., 2007;Ceriani et al., 2019;Holman et al., 2019), it remains robust and constant in zebrafish lateral-line organs, even after the system is mature (***Figure 1e-e’***). It is possible activity in mammalian supporting cells may still be present after maturation but that the activity patterns take a different form. This idea is consistent with work in mice demonstrating that there are periodic waves of calcium in supporting cells in the mature auditory epithelium–distinct from calcium signals that occur during development (Sirko et al., 2019). In addition, studies have shown that during damage, ATP-dependent calcium signals can be induced in the mouse auditory epithelium (Lahne and Gale, 2008). Whether related calcium signals are present in lateral-line supporting cells awaits future work.

The retention of spontaneous activity in supporting cells in the zebrafish lateral line after maturation could also be related to another difference between hair cell systems in zebrafish (and other nonmammalian vertebrates) and mammals–the ability to regenerate hair cells (Corwin and Warchol, 1991;Atkinson et al., 2015;Thomas et al., 2015;Kniss et al., 2016). For example, in zebrafish, after hair-cell loss, supporting cells proliferate to give rise to new hair cells–a process that happens rarely in mature hair cell systems in mammals (Balak et al., 1990;Lush et al., 2019;Thomas and Raible, 2019). The retention of spontaneous calcium signals in zebrafish supporting cells may be important to trigger gene expression required for cell proliferation during regeneration. In this scenario, the retention of spontaneous activity in supporting cells could reflect a feature of their innate regenerative capacity.

Although supporting cells may be less active in mammals upon maturation, it is likely that they play a vital role akin to glia in the nervous system. Like supporting cells, in the nervous system, Gq-coupled GPCRs such as P2YR1 trigger ER calcium release in glia (Parpura et al., 2011;Shigetomi et al., 2018). In the nervous system these calcium signals can be modulated or synchronized with neuronal activity (Aguado et al., 2002), and are linked to several pathological conditions including: seizure initiation in epilepsy, detection of cellular death as well as excitotoxic neurotransmission (Ding et al., 2007;Parpura et al., 2011;Pascual et al., 2012;Alves et al., 2017). In the future it will be important to study the role supporting cells play in these contexts in hair cell systems. Given the conservation of calcium signal generation in supporting cells, the lateral-line system provides an excellent model to study these signals *in vivo* in future work.

### The origin and role of spontaneous calcium activity in zebrafish cholinergic efferents

Our work demonstrates that cholinergic efferents in the lateral-line system are spontaneously active (***Figure 3a1-a2’, Figure 3—figure supplement 1a1-a2’***). In addition, spontaneous activity in cholinergic efferents is stronger during development, consistent with a vital developmental role (***Figure 3—figure supplement 1c-c’***). Numerous studies have investigated the role of cholinergic efferents in mature hair cell systems. In this context, cholinergic efferents are inhibitory, either through their contacts with hair cells (outer auditory hair cells, type II vestibular hair cells, lateral-line hair cells) or afferents terminals (inner auditory hair cells, type I vestibular hair cells) (Housley and Ashmore, 1991;Fuchs and Murrow, 1992;Simmons, 2002;Poppi et al., 2020). But what role these efferents play during development is less clear. Work on auditory inner hair cells in mice has shown that cholinergic efferents are important for synapse maturation–if efferent function is impaired hair cell synapses display defects in calcium-vesicle coupling (Johnson et al., 2013). In the developing vestibular system, it has also been shown that application of acetylcholine can evoke and modulate calcium increases in developing type 1 hair cells and afferent neurons (Holman et al., 2019). Our study suggests the activity of cholinergic efferents does not directly influence or trigger spontaneous calcium activity in immature lateral-line hair cells (***Figure 3c-c’***). It is possible that under the native conditions present within our system, the amount of acetylcholine released from the efferent fibers is not sufficient to alter hair cell calcium to a level detectable using GECIs. It is also possible that acetylcholine may act on supporting cells, other efferent neurons, or afferents neurons in the developing lateral line. Recent work in the mouse vestibular system has shown that application of acetylcholine can evoke calcium responses in supporting cells (Rabbitt and Holman, 2021). In the future it will be important examine the impact and target of cholinergic efferent neurons in the lateral line more closely, in the context of development.

In addition to studies focused on the periphery, work in mice has shown that hair cell spontaneous activity is mirrored downstream, in the central auditory pathway (Babola et al., 2018). Furthermore, spontaneous activity from each ear is patterned centrally, in a bilateral manner. Recent work has shown that cholinergic efferent neurons are required to ensure that spontaneous activity is patterned in a bilateral manner (Babola et al., 2021;Wang et al., 2021). In our work, we demonstrated that spontaneous activity is highly correlated at all efferent terminals within a neuromast. We also found that correlated activity in the cholinergic efferent (likely the same fiber) extends to multiple neuromasts along one side of the fish body (***Figure 3—figure supplement 2a-b***). Although spontaneous activity in the cholinergic efferents terminals does not positively or negatively correlate with hair cell spontaneous activity (***Figure 3b***), it is possible that these efferents may regulate hair cell or afferent spontaneous activity in a manner that is bilateral (left versus right side of the fish body). In the future it will be interesting to develop methods to image activity from cholinergic efferents innervating each side of the zebrafish body. In addition, it will be interesting to determine how hair cell spontaneous activity is reflected in the brain, and whether cholinergic efferents impact the patterning of this activity more centrally.

Could cholinergic efferents pattern bilaterality? Early studies in fishes demonstrated that cholinergic efferents suppresses afferent neuron activity during self-generated movements (Russell, 1968;Roberts and Russell, 1972;Russell and Roberts, 1974). More recent work has shown in mature zebrafish larvae, during swimming cholinergic efferents are synchronously active with spinal motor neurons–this synchronized activity can inhibit hair cell afferents (Lunsford et al., 2019;Pichler and Lagnado, 2020). Therefore, one possibility is that during development, spontaneous activity in cholinergic efferents may also reflect the activity of spinal motor neurons. During spinal cord development, the central pattern generator (CPG) is established, in order to create the oscillatory rhythms needed for locomotion (Grillner et al., 1998;Myers et al., 2005). As part of CPG maturation, a rhythmic alternation between the two sides of the spinal cord is established. This motor prerequisite to swimming behavior may also synchronize with spontaneous activity in cholinergic efferents of the lateral line. If the cholinergic efferents set up bilateral patterning of the lateral line, tapping into the activity of the developing spinal motor neurons and the CPG is a viable way for this to occur (***Figure 7***).

In summary our work provides a comprehensive characterization of spontaneous activity in the zebrafish lateral line. Using LSFM we demonstrate that in neuromast organs, spontaneous activity is present in hair cells, supporting cells and cholinergic efferent terminals. Within the lateral-line, each of these activities may play unique roles in sensory system function, development, maintenance, regeneration, and response to damage. Exploring the role of these spontaneous activities in these important biological contexts will make for exciting future work.

## Supporting information

Movie 1_HC_3D_cytoGCaMP

Movie 2_HC_2D_cytoGCaMP

Movie 3_SC_cytoGCaMP

Movie 4_HC_SC_2_color

Movie 5_HC_eff_2_color

Movie 6_HC_bundle_base_memGCaMP

## Acknowledgements

We acknowledge Alisha Beirl for helping to make the *she:GCaMP6s* and *myo6b:GCaMP6s* transgenic lines. We thank Nathan Lawson and Tatjana Piotrowski for providing the *she* promoter and Jennifer Li for sending the *Gal4:Chat;UAS:GCaMP6s* transgenic line. We would also like to thank Osama Hamdi for his contribution to early Matlab analyses of spontaneous calcium activities. We thank Lavinia Sheets, Cat Weisz and Juan Angueyra for thoughtful comments on the manuscript.

## Data availability

Data that support the findings of this study are available from the corresponding author upon reasonable request.

## Code availability

Matlab R2020a used to process functional imaging data and the code is available upon request.

## References

Ackman, J.B., Burbridge, T.J., and Crair, M.C. (2012). Retinal waves coordinate patterned activity throughout the developing visual system. Nature 490, 219–225.

Aguado, F., Espinosa-Parrilla, J.F., Carmona, M.A., and Soriano, E. (2002). Neuronal activity regulates correlated network properties of spontaneous calcium transients in astrocytes in situ. J Neurosci 22, 9430–9444.

Akrouh, A., and Kerschensteiner, D. (2013). Intersecting circuits generate precisely patterned retinal waves. Neuron 79, 322–334.

Allene, C., Cattani, A., Ackman, J.B., Bonifazi, P., Aniksztejn, L., Ben-Ari, Y., and Cossart, R. (2008). Sequential generation of two distinct synapse-driven network patterns in developing neocortex. Journal of Neuroscience 28, 12851–12863.

Alves, M., Gomez-Villafuertes, R., Delanty, N., Farrell, M.A., O’brien, D.F., Miras-Portugal, M.T., Hernandez, M.D., Henshall, D.C., and Engel, T. (2017). Expression and function of the metabotropic purinergic P2Y receptor family in experimental seizure models and patients with drug-refractory epilepsy. Epilepsia 58, 1603–1614.

Arganda-Carreras, I., Sorzano, C.O., Marabini, R., Carazo, J.M., Ortiz-De-Solorzano, C., and Kybic, J. (Year). “Consistent and elastic registration of histological sections using vector-spline regularization”, in: International Workshop on Computer Vision Approaches to Medical Image Analysis: Springer), 85–95.

Aschoff, A., and Ostwald, J. (1987). Different origins of cochlear efferents in some bat species, rats, and guinea pigs. J Comp Neurol 264, 56–72.

Assad, J.A., Shepherd, G.M., and Corey, D.P. (1991). Tip-link integrity and mechanical transduction in vertebrate hair cells. Neuron 7, 985–994.

Atkinson, P.J., Huarcaya Najarro, E., Sayyid, Z.N., and Cheng, A.G. (2015). Sensory hair cell development and regeneration: similarities and differences. Development 142, 1561–1571.

Babola, T.A., Kersbergen, C.J., Wang, H.C., and Bergles, D.E. (2020). Purinergic signaling in cochlear supporting cells reduces hair cell excitability by increasing the extracellular space. Elife 9, e52160.

Babola, T.A., Li, S., Gribizis, A., Lee, B.J., Issa, J.B., Wang, H.C., Crair, M.C., and Bergles, D.E. (2018). Homeostatic control of spontaneous activity in the developing auditory system. Neuron 99, 511–524. e515.

Babola, T.A., Li, S., Wang, Z., Kersbergen, C.J., Elgoyhen, A.B., Coate, T.M., and Bergles, D.E. (2021). Purinergic signaling controls spontaneous activity in the auditory system throughout early development. Journal of Neuroscience 41, 594–612.

Balak, K.J., Corwin, J.T., and Jones, J.E. (1990). Regenerated hair cells can originate from supporting cell progeny: evidence from phototoxicity and laser ablation experiments in the lateral line system. J Neurosci 10, 2502–2512.

Bansal, A., Singer, J.H., Hwang, B.J., Xu, W., Beaudet, A., and Feller, M.B. (2000). Mice lacking specific nicotinic acetylcholine receptor subunits exhibit dramatically altered spontaneous activity patterns and reveal a limited role for retinal waves in forming ON and OFF circuits in the inner retina. Journal of Neuroscience 20, 7672–7681.

Bechstedt, S., and Howard, J. (2007). Models of hair cell mechanotransduction. Current topics in membranes 59, 399–424.

Brandt, A., Striessnig, J., and Moser, T. (2003). CaV1.3 channels are essential for development and presynaptic activity of cochlear inner hair cells. J Neurosci 23, 10832–10840.

Bucheimer, R.E., and Linden, J. (2004). Purinergic regulation of epithelial transport. The Journal of physiology 555, 311–321.

Carpaneto Freixas, A.E., Moglie, M.J., Castagnola, T., Salatino, L., Domene, S., Marcovich, I., Gallino, S., Wedemeyer, C., Goutman, J.D., Plazas, P.V., and Elgoyhen, A.B. (2021). Unraveling the Molecular Players at the Cholinergic Efferent Synapse of the Zebrafish Lateral Line. J Neurosci 41, 47–60.

Ceriani, F., Hendry, A., Jeng, J.Y., Johnson, S.L., Stephani, F., Olt, J., Holley, M.C., Mammano, F., Engel, J., Kros, C.J., Simmons, D.D., and Marcotti, W. (2019). Coordinated calcium signalling in cochlear sensory and non-sensory cells refines afferent innervation of outer hair cells. EMBO J 38.

Clause, A., Kim, G., Sonntag, M., Weisz, C.J., Vetter, D.E., Rűbsamen, R., and Kandler, K. (2014). The precise temporal pattern of prehearing spontaneous activity is necessary for tonotopic map refinement. Neuron 82, 822–835.

Corns, L.F., Johnson, S.L., Roberts, T., Ranatunga, K.M., Hendry, A., Ceriani, F., Safieddine, S., Steel, K.P., Forge, A., Petit, C., Furness, D.N., Kros, C.J., and Marcotti, W. (2018). Mechanotransduction is required for establishing and maintaining mature inner hair cells and regulating efferent innervation. Nat Commun 9, 4015.

Corwin, J.T., and Warchol, M.E. (1991). Auditory hair cells: structure, function, development, and regeneration. Annual review of neuroscience 14, 301–333.

Courjaret, R., Dib, M., and Machaca, K. (2018). Spatially restricted subcellular Ca(2+) signaling downstream of store-operated calcium entry encoded by a cortical tunneling mechanism. Sci Rep 8, 11214.

Desai, B.N., and Leitinger, N. (2014). Purinergic and calcium signaling in macrophage function and plasticity. Frontiers in immunology 5, 580.

Ding, S., Fellin, T., Zhu, Y., Lee, S.Y., Auberson, Y.P., Meaney, D.F., Coulter, D.A., Carmignoto, G., and Haydon, P.G. (2007). Enhanced astrocytic Ca2+ signals contribute to neuronal excitotoxicity after status epilepticus. J Neurosci 27, 10674–10684.

Dow, E., Jacobo, A., Hossain, S., Siletti, K., and Hudspeth, A.J. (2018). Connectomics of the zebrafish’s lateral-line neuromast reveals wiring and miswiring in a simple microcircuit. Elife 7, e33988.

Dulon, D., Moataz, R., and Mollard, P. (1993). Characterization of Ca2+ signals generated by extracellular nucleotides in supporting cells of the organ of Corti. Cell Calcium 14, 245–254.

Eckrich, T., Blum, K., Milenkovic, I., and Engel, J. (2018). Fast Ca2+ transients of inner hair cells arise coupled and uncoupled to Ca2+ waves of inner supporting cells in the developing mouse cochlea. Frontiers in molecular neuroscience 11, 264.

Edelstein, A., Amodaj, N., Hoover, K., Vale, R., and Stuurman, N. (2010). Computer control of microscopes using microManager. Curr Protoc Mol Biol Chapter 14, Unit14 20.

Förster, D., Arnold-Ammer, I., Laurell, E., Barker, A.J., Fernandes, A.M., Finger-Baier, K., Filosa, A., Helmbrecht, T.O., Kölsch, Y., and Kühn, E. (2017). Genetic targeting and anatomical registration of neuronal populations in the zebrafish brain with a new set of BAC transgenic tools. Scientific reports 7, 1–11.

Freixas, A.E.C., Moglie, M.J., Castagnola, T., Salatino, L., Domene, S., Marcovich, I., Gallino, S., Wedemeyer, C., Goutman, J.D., and Plazas, P.V. (2021). Unraveling the Molecular Players at the Cholinergic Efferent Synapse of the Zebrafish Lateral Line. Journal of Neuroscience 41, 47–60.

Fuchs, P.A., and Murrow, B.W. (1992). A Novel Cholinergic Receptor Mediates Inhibition of Chick Cochlear Hair-Cells. Proceedings of the Royal Society B-Biological Sciences 248, 35–40.

Glowatzki, E., and Fuchs, P.A. (2000). Cholinergic synaptic inhibition of inner hair cells in the neonatal mammalian cochlea. Science 288, 2366–2368.

Goodyear, R.J., and Richardson, G.P. (2003). A novel antigen sensitive to calcium chelation that is associated with the tip links and kinocilial links of sensory hair bundles. Journal of Neuroscience 23, 4878–4887.

Grillner, S., Ekeberg, El Manira, A., Lansner, A., Parker, D., Tegner, J., and Wallen, P. (1998). Intrinsic function of a neuronal network - a vertebrate central pattern generator. Brain Res Brain Res Rev 26, 184–197.

Guo, M., Li, Y., Su, Y., Lambert, T., Nogare, D.D., Moyle, M.W., Duncan, L.H., Ikegami, R., Santella, A., Rey-Suarez, I., Green, D., Beiriger, A., Chen, J., Vishwasrao, H., Ganesan, S., Prince, V., Waters, J.C., Annunziata, C.M., Hafner, M., Mohler, W.A., Chitnis, A.B., Upadhyaya, A., Usdin, T.B., Bao, Z., Colon-Ramos, D., La Riviere, P., Liu, H., Wu, Y., and Shroff, H. (2020). Rapid image deconvolution and multiview fusion for optical microscopy. Nat Biotechnol 38, 1337–1346.

Hiel, H., Luebke, A.E., and Fuchs, P.A. (2000). Cloning and expression of the alpha9 nicotinic acetylcholine receptor subunit in cochlear hair cells of the chick. Brain Res 858, 215–225.

Hiu-Tung, C.W., Zhang, Q., Beirl, A.J., Petralia, R.S., Wang, Y.-X., and Kindt, K. (2019). Synaptic mitochondria regulate hair-cell synapse size and function. Elife 8, e48914.

Holman, H.A., Poppi, L.A., Frerck, M., and Rabbitt, R.D. (2019). Spontaneous and acetylcholine evoked calcium transients in the developing mouse utricle. Frontiers in cellular neuroscience 13, 186.

Housley, G.D., and Ashmore, J.F. (1991). Direct Measurement of the Action of Acetylcholine on Isolated Outer Hair-Cells of the Guinea-Pig Cochlea. Proceedings of the Royal Society B-Biological Sciences 244, 161–167.

Huang, C.-H., and Moser, T. (2018). Ca2+ regulates the kinetics of synaptic vesicle fusion at the afferent inner hair cell synapse. Frontiers in cellular neuroscience 12, 364.

Huberman, A.D., Feller, M.B., and Chapman, B. (2008). Mechanisms underlying development of visual maps and receptive fields. Annu. Rev. Neurosci. 31, 479–509.

Jeng, J.Y., Ceriani, F., Hendry, A., Johnson, S.L., Yen, P., Simmons, D.D., Kros, C.J., and Marcotti, W. (2020). Hair cell maturation is differentially regulated along the tonotopic axis of the mammalian cochlea. Journal of Physiology-London 598, 151–170.

Jiang, T., Kindt, K., and Wu, D.K. (2017). Transcription factor Emx2 controls stereociliary bundle orientation of sensory hair cells. Elife 6, e23661.

Johnson, S.L., Eckrich, T., Kuhn, S., Zampini, V., Franz, C., Ranatunga, K.M., Roberts, T.P., Masetto, S., Knipper, M., and Kros, C.J. (2011). Position-dependent patterning of spontaneous action potentials in immature cochlear inner hair cells. Nature neuroscience 14, 711–717.

Johnson, S.L., Kennedy, H.J., Holley, M.C., Fettiplace, R., and Marcotti, W. (2012). The resting transducer current drives spontaneous activity in prehearing mammalian cochlear inner hair cells. Journal of Neuroscience 32, 10479–10483.

Johnson, S.L., Wedemeyer, C., Vetter, D.E., Adachi, R., Holley, M.C., Elgoyhen, A.B., and Marcotti, W. (2013). Cholinergic efferent synaptic transmission regulates the maturation of auditory hair cell ribbon synapses. Open Biol 3, 130163.

Katz, E., Elgoyhen, A.B., Gomez-Casati, M.E., Knipper, M., Vetter, D.E., Fuchs, P.A., and Glowatzki, E. (2004). Developmental regulation of nicotinic synapses on cochlear inner hair cells. J Neurosci 24, 7814–7820.

Khazipov, R., Sirota, A., Leinekugel, X., Holmes, G.L., Ben-Ari, Y., and Buzsáki, G. (2004). Early motor activity drives spindle bursts in the developing somatosensory cortex. Nature 432, 758–761.

Kikuchi, T., Kimura, R.S., Paul, D.L., Takasaka, T., and Adams, J.C. (2000). Gap junction systems in the mammalian cochlea. Brain research reviews 32, 163–166.

Kindt, K.S., Finch, G., and Nicolson, T. (2012). Kinocilia mediate mechanosensitivity in developing zebrafish hair cells. Developmental cell 23, 329–341.

Kniss, J.S., Jiang, L., and Piotrowski, T. (2016). Insights into sensory hair cell regeneration from the zebrafish lateral line. Current opinion in genetics & development 40, 32–40.

Kros, C.J., Ruppersberg, J.P., and Rusch, A. (1998). Expression of a potassium current in inner hair cells during development of hearing in mice. Nature 394, 281–284.

Kumar, A., Wu, Y., Christensen, R., Chandris, P., Gandler, W., Mccreedy, E., Bokinsky, A., Colón-Ramos, D.A., Bao, Z., and Mcauliffe, M. (2014). Dual-view plane illumination microscopy for rapid and spatially isotropic imaging. Nature protocols 9, 2555–2573.

Kwan, K.M., Fujimoto, E., Grabher, C., Mangum, B.D., Hardy, M.E., Campbell, D.S., Parant, J.M., Yost, H.J., Kanki, J.P., and Chien, C.B. (2007). The Tol2kit: a multisite gateway-based construction kit for Tol2 transposon transgenesis constructs. Developmental dynamics: an official publication of the American Association of Anatomists 236, 3088–3099.

Lahne, M., and Gale, J.E. (2008). Damage-induced activation of ERK1/2 in cochlear supporting cells is a hair cell death-promoting signal that depends on extracellular ATP and calcium. J Neurosci 28, 4918–4928.

Lautermann, J., Wouter-Jan, F., Altenhoff, P., Grümmer, R., Traub, O., Frank, H.-G., Jahnke, K., and Winterhager, E. (1998). Expression of the gap-junction connexins 26 and 30 in the rat cochlea. Cell and tissue research 294, 415–420.

Leighton, A.H., and Lohmann, C. (2016). The wiring of developing sensory circuits—from patterned spontaneous activity to synaptic plasticity mechanisms. Frontiers in neural circuits 10, 71.

Levic, S., Lv, P., and Yamoah, E.N. (2011). The Activity of Spontaneous Action Potentials in Developing Hair Cells Is Regulated by Ca2+-Dependence of a Transient K+ Current. Plos one 6, e29005.

Lobas, M.A., Tao, R., Nagai, J., Kronschlager, M.T., Borden, P.M., Marvin, J.S., Looger, L.L., and Khakh, B.S. (2019). A genetically encoded single-wavelength sensor for imaging cytosolic and cell surface ATP. Nat Commun 10, 711.

Lukasz, D., and Kindt, K.S. (2018). In Vivo Calcium Imaging of Lateral-line Hair Cells in Larval Zebrafish. J Vis Exp.

Lunsford, E.T., Skandalis, D.A., and Liao, J.C. (2019). Efferent modulation of spontaneous lateral line activity during and after zebrafish motor commands. J Neurophysiol 122, 2438–2448.

Lush, M.E., Diaz, D.C., Koenecke, N., Baek, S., Boldt, H., St Peter, M.K., Gaitan-Escudero, T., Romero-Carvajal, A., Busch-Nentwich, E.M., Perera, A.G., Hall, K.E., Peak, A., Haug, J.S., and Piotrowski, T. (2019). scRNA-Seq reveals distinct stem cell populations that drive hair cell regeneration after loss of Fgf and Notch signaling. Elife 8.

Marcotti, W., Erven, A., Johnson, S.L., Steel, K.P., and Kros, C.J. (2006). Tmc1 is necessary for normal functional maturation and survival of inner and outer hair cells in the mouse cochlea. J Physiol 574, 677–698.

Marcotti, W., Johnson, S.L., Rusch, A., and Kros, C.J. (2003). Sodium and calcium currents shape action potentials in immature mouse inner hair cells. J Physiol 552, 743–761.

Moody, W.J., and Bosma, M.M. (2005). Ion channel development, spontaneous activity, and activity-dependent development in nerve and muscle cells. Physiological reviews 85, 883–941.

Müller, U. (2008). Cadherins and mechanotransduction by hair cells. Current opinion in cell biology 20, 557–566.

Mulroy, M.J., Dempewolf, S.A., Curtis, S., and Iida, H.C. (1993). Gap junctional connections between hair cells, supporting cells and nerves in a vestibular organ. Hearing research 71, 98–105.

Myers, C.P., Lewcock, J.W., Hanson, M.G., Gosgnach, S., Aimone, J.B., Gage, F.H., Lee, K.F., Landmesser, L.T., and Pfaff, S.L. (2005). Cholinergic input is required during embryonic development to mediate proper assembly of spinal locomotor circuits. Neuron 46, 37–49.

Newbold, D.J., Laumann, T.O., Hoyt, C.R., Hampton, J.M., Montez, D.F., Raut, R.V., Ortega, M., Mitra, A., Nielsen, A.N., and Miller, D.B. (2020). Plasticity and spontaneous activity pulses in disused human brain circuits. Neuron 107, 580–589. e586.

Obholzer, N., Wolfson, S., Trapani, J.G., Mo, W., Nechiporuk, A., Busch-Nentwich, E., Seiler, C., Sidi, S., Sollner, C., Duncan, R.N., Boehland, A., and Nicolson, T. (2008). Vesicular glutamate transporter 3 is required for synaptic transmission in zebrafish hair cells. J Neurosci 28, 2110–2118.

Parks, X.X., Contini, D., Jordan, P.M., and Holt, J.C. (2017). Confirming a Role for alpha9nAChRs and SK Potassium Channels in Type II Hair Cells of the Turtle Posterior Crista. Front Cell Neurosci 11, 356.

Parpura, V., Grubisic, V., and Verkhratsky, A. (2011). Ca(2+) sources for the exocytotic release of glutamate from astrocytes. Biochim Biophys Acta 1813, 984–991.

Parslow, A., Cardona, A., and Bryson-Richardson, R.J. (2014). Sample drift correction following 4D confocal time-lapse imaging. Journal of visualized experiments: JoVE.

Pascual, O., Ben Achour, S., Rostaing, P., Triller, A., and Bessis, A. (2012). Microglia activation triggers astrocyte-mediated modulation of excitatory neurotransmission. Proc Natl Acad Sci U S A 109, E197–205.

Peloggia, J., Munch, D., Meneses-Giles, P., Romero-Carvajal, A., Lush, M.E., Lawson, N.D., Mcclain, M., Pan, Y.A., and Piotrowski, T. (2021). Adaptive cell invasion maintains lateral line organ homeostasis in response to environmental changes. Dev Cell 56, 1296–1312 e1297.

Pichler, P., and Lagnado, L. (2020). Motor Behavior Selectively Inhibits Hair Cells Activated by Forward Motion in the Lateral Line of Zebrafish. Curr Biol 30, 150–157 e153.

Pickles, J., Comis, S., and Osborne, M. (1984). Cross-links between stereocilia in the guinea pig organ of Corti, and their possible relation to sensory transduction. Hearing research 15, 103–112.

Poppi, L.A., Holt, J.C., Lim, R., and Brichta, A.M. (2020). A review of efferent cholinergic synaptic transmission in the vestibular periphery and its functional implications. J Neurophysiol 123, 608–629.

Power, R.M., and Huisken, J. (2017). A guide to light-sheet fluorescence microscopy for multiscale imaging. Nat Methods 14, 360–373.

Quillien, A., Abdalla, M., Yu, J., Ou, J., Zhu, L.J., and Lawson, N.D. (2017). Robust Identification of Developmentally Active Endothelial Enhancers in Zebrafish Using FANS-Assisted ATAC-Seq. Cell Rep 20, 709–720.

Quirin, S., Vladimirov, N., Yang, C.T., Peterka, D.S., Yuste, R., and Ahrens, M.B. (2016). Calcium imaging of neural circuits with extended depth-of-field light-sheet microscopy. Opt Lett 41, 855–858.

Rabbitt, R.D., and Holman, H.A. (2021). ATP and ACh Evoked Calcium Transients in the Neonatal Mouse Cochlear and Vestibular Sensory Epithelia. Frontiers in Neuroscience, 1175.

Raible, D.W., and Kruse, G.J. (2000). Organization of the lateral line system in embryonic zebrafish. J Comp Neurol 421, 189–198.

Roberts, B.L., and Russell, I.J. (1972). The activity of lateral-line efferent neurones in stationary and swimming dogfish. J Exp Biol 57, 435–448.

Roux, I., Wersinger, E., Mcintosh, J.M., Fuchs, P.A., and Glowatzki, E. (2011). Onset of cholinergic efferent synaptic function in sensory hair cells of the rat cochlea. Journal of Neuroscience 31, 15092–15101.

Russell, I.J. (1968). Influence of efferent fibres on a receptor. Nature 219, 177–178.

Russell, I.J., and Roberts, B.L. (1974). Active Reduction of Lateral-Line Sensitivity in Swimming Dogfish. Journal of Comparative Physiology 94, 7–15.

Sato, Y., Handa, T., Matsumura, M., and Orita, Y. (1998). Gap junction change in supporting cells of the organ of Corti with ryanodine and caffeine. Acta oto-laryngologica 118, 821–825.

Schneider, C.A., Rasband, W.S., and Eliceiri, K.W. (2012). NIH Image to ImageJ: 25 years of image analysis. Nat Methods 9, 671–675.

Seiler, C., Finger-Baier, K.C., Rinner, O., Makhankov, Y.V., Schwarz, H., Neuhauss, S.C., and Nicolson, T. (2005). Duplicated genes with split functions: independent roles of protocadherin15 orthologues in zebrafish hearing and vision. Development 132, 615–623.

Sendin, G., Bourien, J., Rassendren, F., Puel, J.-L., and Nouvian, R. (2014). Spatiotemporal pattern of action potential firing in developing inner hair cells of the mouse cochlea. Proceedings of the National Academy of Sciences 111, 1999–2004.

Sheets, L., He, X.J., Olt, J., Schreck, M., Petralia, R.S., Wang, Y.X., Zhang, Q., Beirl, A., Nicolson, T., Marcotti, W., Trapani, J.G., and Kindt, K.S. (2017). Enlargement of ribbons in zebrafish hair cells increases calcium currents, but disrupts afferent spontaneous activity and timing of stimulus onset. J Neurosci.

Sheets, L., Kindt, K.S., and Nicolson, T. (2012). Presynaptic CaV1. 3 channels regulate synaptic ribbon size and are required for synaptic maintenance in sensory hair cells. Journal of Neuroscience 32, 17273–17286.

Shigetomi, E., Hirayama, Y.J., Ikenaka, K., Tanaka, K.F., and Koizumi, S. (2018). Role of Purinergic Receptor P2Y1 in Spatiotemporal Ca(2+) Dynamics in Astrocytes. J Neurosci 38, 1383–1395.

Sidi, S., Busch-Nentwich, E., Friedrich, R., Schoenberger, U., and Nicolson, T. (2004). gemini encodes a zebrafish L-type calcium channel that localizes at sensory hair cell ribbon synapses. J Neurosci 24, 4213–4223.

Simmons, D.D. (2002). Development of the inner ear efferent system across vertebrate species. J Neurobiol 53, 228–250.

Sirko, P., Gale, J.E., and Ashmore, J.F. (2019). Intercellular Ca(2+) signalling in the adult mouse cochlea. J Physiol 597, 303–317.

Suli, A., Watson, G.M., Rubel, E.W., and Raible, D.W. (2012). Rheotaxis in larval zebrafish is mediated by lateral line mechanosensory hair cells. PLoS One 7, e29727.

Sun, S., Babola, T., Pregernig, G., So, K.S., Nguyen, M., Su, S.M., Palermo, A.T., Bergles, D.E., Burns, J.C., and Muller, U. (2018). Hair Cell Mechanotransduction Regulates Spontaneous Activity and Spiral Ganglion Subtype Specification in the Auditory System. Cell 174, 1247–1263 e1215.

Thevenaz, P., Ruttimann, U.E., and Unser, M. (1998). A pyramid approach to subpixel registration based on intensity. IEEE transactions on image processing 7, 27–41.

Thomas, E.D., Cruz, I.A., Hailey, D.W., and Raible, D.W. (2015). There and back again: development and regeneration of the zebrafish lateral line system. Wiley Interdisciplinary Reviews: Developmental Biology 4, 1–16.

Thomas, E.D., and Raible, D.W. (2019). Distinct progenitor populations mediate regeneration in the zebrafish lateral line. Elife 8.

Trapani, J.G., and Nicolson, T. (2011). Mechanism of spontaneous activity in afferent neurons of the zebrafish lateral-line organ. J Neurosci 31, 1614–1623.

Tritsch, N.X., Yi, E., Gale, J.E., Glowatzki, E., and Bergles, D.E. (2007). The origin of spontaneous activity in the developing auditory system. Nature 450, 50–55.

Vladimirov, N., Mu, Y., Kawashima, T., Bennett, D.V., Yang, C.T., Looger, L.L., Keller, P.J., Freeman, J., and Ahrens, M.B. (2014). Light-sheet functional imaging in fictively behaving zebrafish. Nat Methods 11, 883–884.

Wang, H.C., Lin, C.-C., Cheung, R., Zhang-Hooks, Y., Agarwal, A., Ellis-Davies, G., Rock, J., and Bergles, D.E. (2015). Spontaneous activity of cochlear hair cells triggered by fluid secretion mechanism in adjacent support cells. Cell 163, 1348–1359.

Wang, Y., Sanghvi, M., Gribizis, A., Zhang, Y., Song, L., Morley, B., Barson, D.G., Santos-Sacchi, J., Navaratnam, D., and Crair, M. (2021). Efferent feedback controls bilateral auditory spontaneous activity. Nature communications 12, 1–16.

Warland, D.K., Huberman, A.D., and Chalupa, L.M. (2006). Dynamics of spontaneous activity in the fetal macaque retina during development of retinogeniculate pathways. Journal of Neuroscience 26, 5190–5197.

Wosniack, M.E., Kirchner, J.H., Chao, L.-Y., Zabouri, N., Lohmann, C., and Gjorgjieva, J. (2021). Adaptation of spontaneous activity in the developing visual cortex. Elife 10, e61619.

Zhang, Q., Li, S., Wong, H.C., He, X.J., Beirl, A., Petralia, R.S., Wang, Y.X., and Kindt, K.S. (2018). Synaptically silent sensory hair cells in zebrafish are recruited after damage. Nat Commun 9, 1388.

Zhang, Q.X., He, X.J., Wong, H.C., and Kindt, K.S. (2016). Functional calcium imaging in zebrafish lateral-line hair cells. Methods Cell Biol 133, 229–252.

