## Supplemental figures and Legends for "Using light-sheet microscopy to study spontaneous activity in the developing lateral-line system"

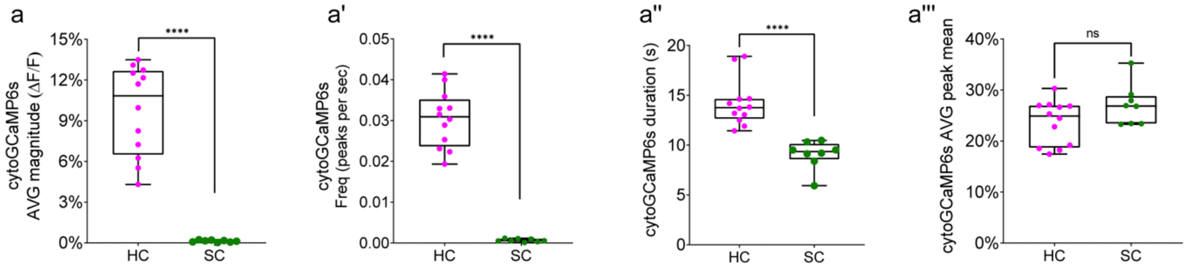


**Figure 1—figure supplement 1. Comparison of spontaneous calcium activities between hair cells and supporting cells.**

(a-a''') Quantification of spontaneous calcium activities in hair cells and supporting cells at day 3 reveals significant differences in the average magnitude (a), frequency (a') and the peak duration (a''). The peak mean is comparable between hair cells and supporting cells (a'''). Each dot in a-a''' represents one neuromast. An unpaired t-test was used in a-a'''. ****p<0.0001.

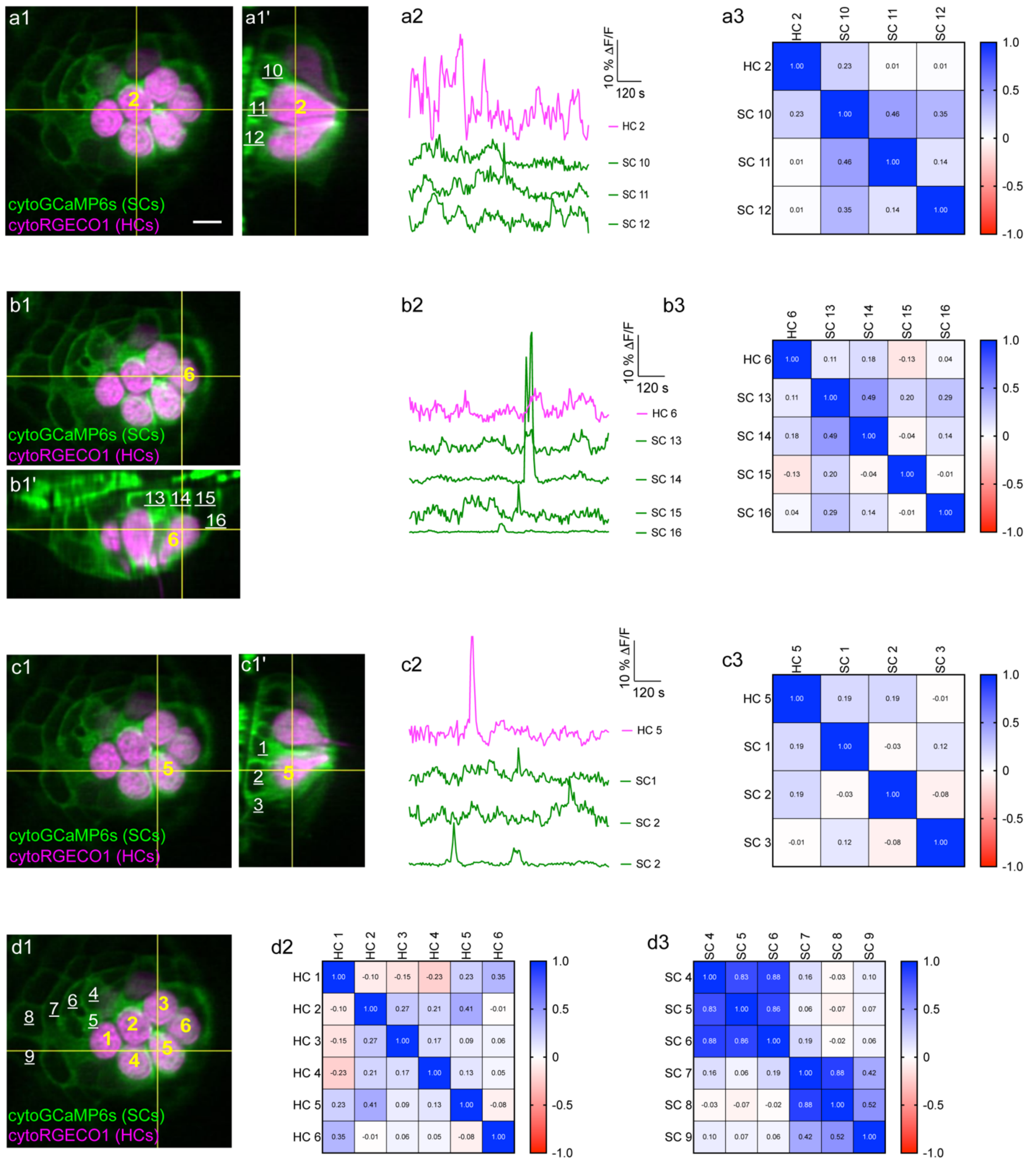


**Figure 1—figure supplement 2. Additional examples of spontaneous calcium activities in hair cells and surrounding supporting cells.** A top-down view (a1) and the corresponding side view (a1') of hair cell 2 from the same day 3 neuromast organ in ***Figure 1***. (a2) The temporal curves of spontaneous calcium activity in hair cell 2 (labeled in a1-a1') and its 3 surrounding supporting cells labeled in a1' show distinct time courses. (a3) Heatmap showing the Pearson's R values between hair cell 2 and the surrounding supporting cells. A top-down view (b1) and the corresponding side view (b1') of hair cell 6 from the same neuromast organ in ***Figure 1***. (b2) The temporal curves of spontaneous calcium activity in hair cell 6 (labeled in b1-b1') and its 4

surrounding supporting cells (labeled in b1') also show distinct time courses between hair cells and the supporting cells. (b3) Heatmap showing the Pearson's R values between hair cell 6 and its surrounding supporting cells. (c1-c3) Additional information from hair cell 5 (labeled in c1-c1') and its surrounding supporting cells (labeled in c1') from the same neuromast organ in ***Figure 1***. (c2) The temporal curves of the cells labeled in c1-c1' and (a3) heatmaps showing the Pearson's R values between hair cell 5 and its surrounding supporting cells. (d1-d3) Heatmaps showing the Pearson's R values from the 6 hair cells (d2) and 6 supporting cells (d3) labeled in d1. Scale bar = 5 µm.

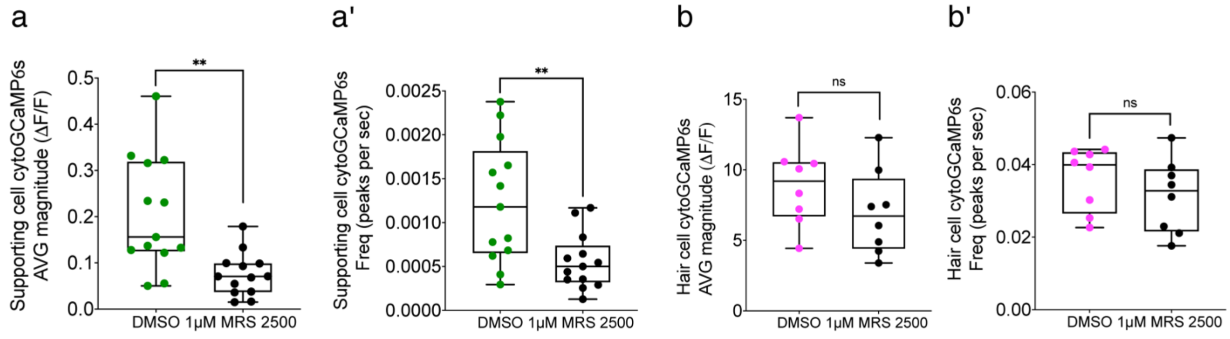


**Figure 2—figure supplement 1. Quantification of P2yr1 block on hair cell and supporting cell spontaneous calcium activities**. Box plots show the average magnitude and frequency of spontaneous calcium activity in the supporting cells (a-a') and hair cells (b-b') before and after 15 min of treatment with 1 µM MRS2500. All measurements were performed in immature neuromasts at day 3. Each point in a-b' represents one neuromast. A paired t-test was used. **p<0.01.

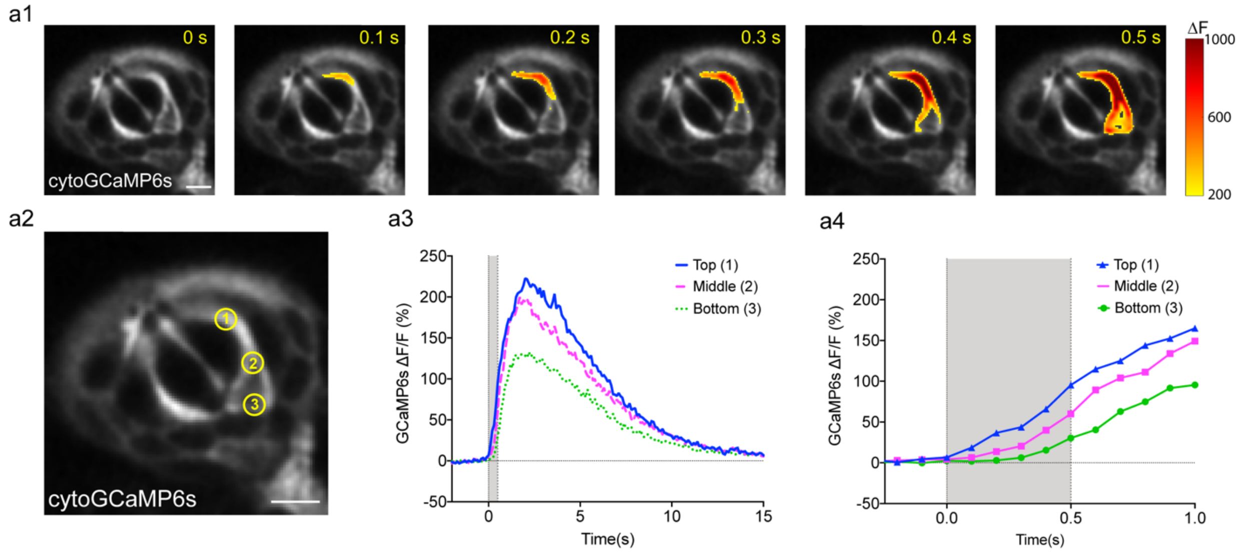


**Figure 2—figure supplement 2. Spontaneous calcium activity in supporting cells initiates in the apex.** (a1) A representative spontaneous event in a supporting cell taken in cross-section view during six 0.1 s time intervals, taken at day 3. Spatial increases in cytoGCaMP6s signal (ΔF) are color coded and indicated by the heatmap. (a2) 3 distinct positions within the responding supporting cell are labeled. (a3) The temporal curve of the calcium signal at each position indicated in a2 during the 18 s window encompassing the spontaneous event. (a4) To better compare the signal onset time, an enlarged profile of the 1.25 s period indicated in the gray bar in a3 is depicted in a4. Measurements were performed at day 3. Scale bars = 5 µm.

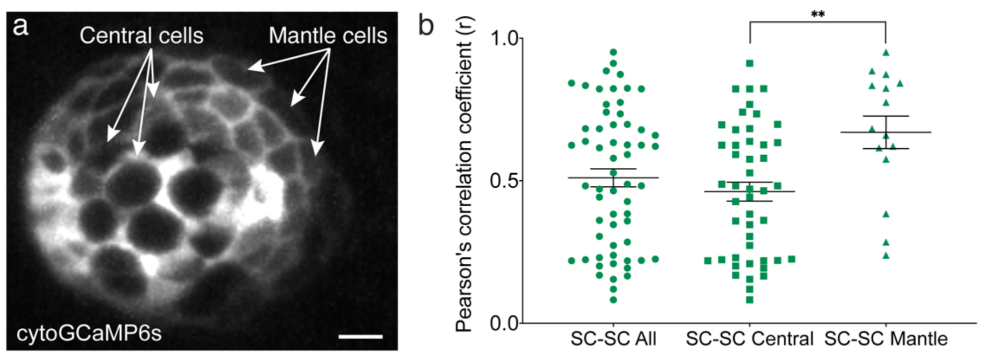


**Figure 2—figure supplement 3. Spontaneous calcium activity in outer supporting cells is more correlated compared to central supporting cells.** (a) An image of supporting cells expressing cytoGCaMP6s at day 3. Examples of the outer-most supporting cells (mantle cells), and central supporting cells are labels with arrows. (b) Pearson's R values comparing the correlation in spontaneous calcium activities between all neighboring supporting cells, neighboring central supporting cells and neighboring outer-most (mantle) supporting cells. Measurements were performed in neuromasts at day 3. A Kruskal-Wallis test was used. **$p<0.01$. Scale bar = 5 µm.

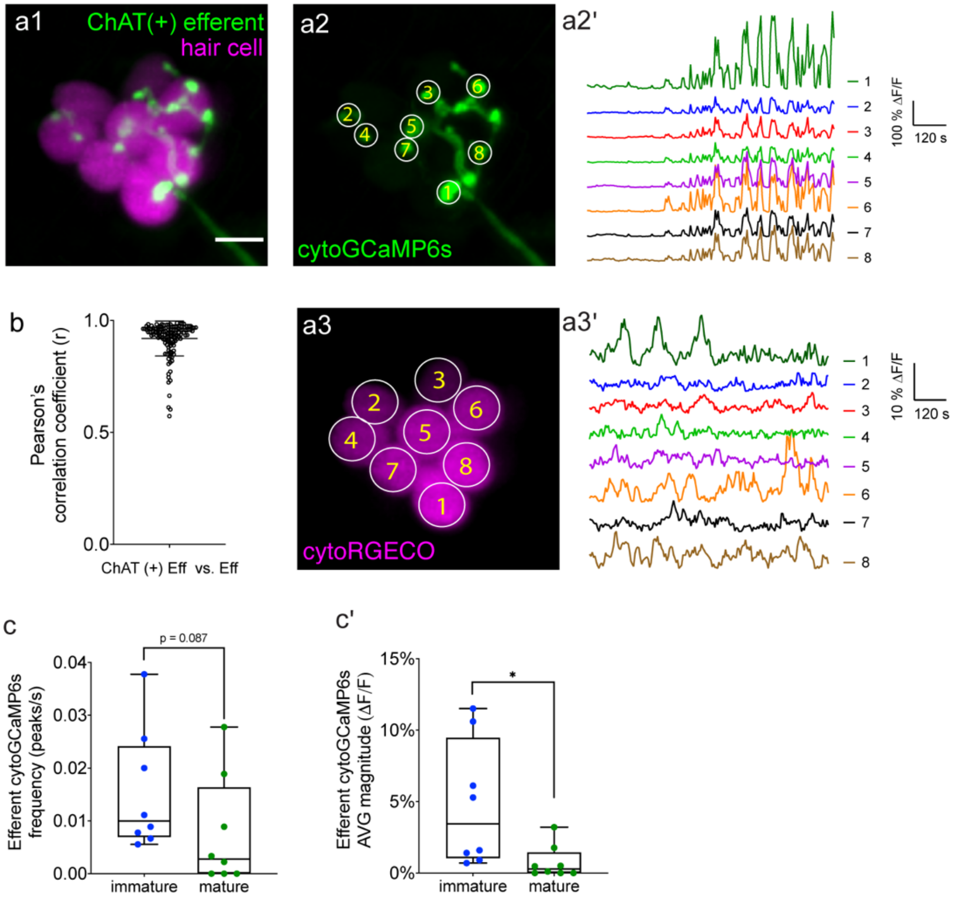


**Figure 3—figure supplement 1. An additional example of two-color imaging of spontaneous activities in hair cells and cholinergic efferent terminals**. (a1) A maximum intensity z-projection image of a double-transgenic zebrafish line expressing the red GECI cytoRGECO1 in hair cells and the green GECI cytoGCaMP6s in the cholinergic efferents at day 3. (a2) The cholinergic efferent terminal image with individual terminals contacting different hair cells in a1 are labeled. (a2') The corresponding temporal curves of spontaneous calcium activities of the 8 efferent terminals indicated in a2. (a3) The individual hair cells are labeled. (a3') The corresponding temporal curves of spontaneous cytosolic calcium activities of those 8 hair cells indicated in a3. (b) Pearson's R values of spontaneous calcium activity between cholinergic efferent terminal in an individual neuromast, n = 129 terminals in 4 neuromasts at day 3. (c-c') Box plots showing the average magnitude of spontaneous calcium activity in cholinergic efferent terminals in immature (day 3) and mature (day 6) neuromasts. Each point in c-c' represents one neuromast. An unpaired t-test was used in c-c'. Scale bar = 5 µm.

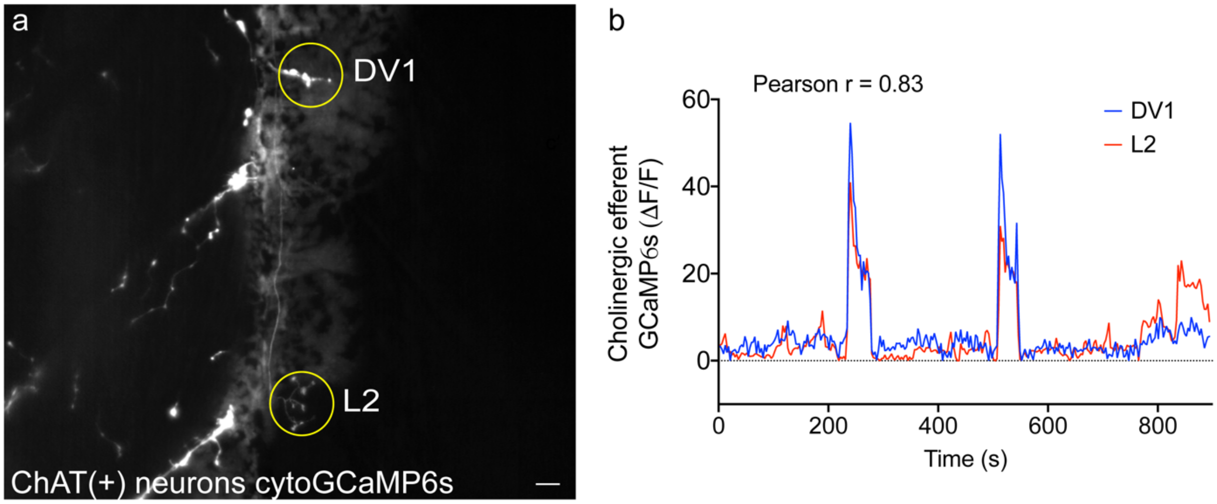


**Figure 3—figure supplement 2. An example of spontaneous activity in cholinergic efferent terminals from two adjacent neuromasts**. (a) A maximum intensity z-projection image of a transgenic zebrafish line expressing cytoGCaMP6s in the cholinergic neurons including the cholinergic efferents terminals of two adjacent neuromasts at day 3 (circled, DV1 and L2) at day 3. (b) The temporal curve of the calcium signal in each circle indicated in (a) during the 900 s time window. The overall spontaneous calcium activities in the cholinergic terminals of these two representative neuromasts is highly correlated. Scale bar = 10 µm.

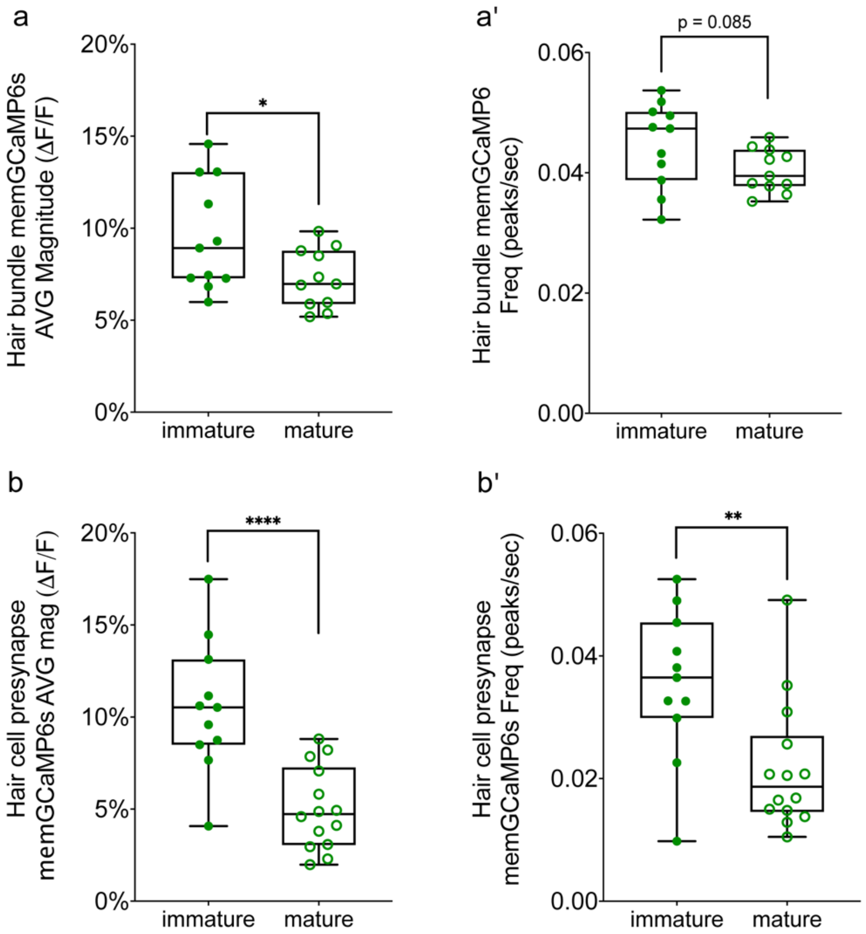


**Figure 4—figure supplement 1. Spontaneous activity in both the hair bundle and presynapse region decrease upon maturation**. Box plots showing the average magnitude (a) and frequency (a') of spontaneous calcium activity at the hair bundle is higher in immature (day 3) versus mature (day 6) hair cells. Similarly, the average magnitude (b) and frequency (b') of spontaneous calcium activity at the presynapse is higher in immature (day 3) versus mature (day 6) hair cells. Each point in a-b' represents one neuromast. An unpaired t-test was used in a-b'. * $p<0.05$, ** $p<0.01$, ****$p<0.0001$.

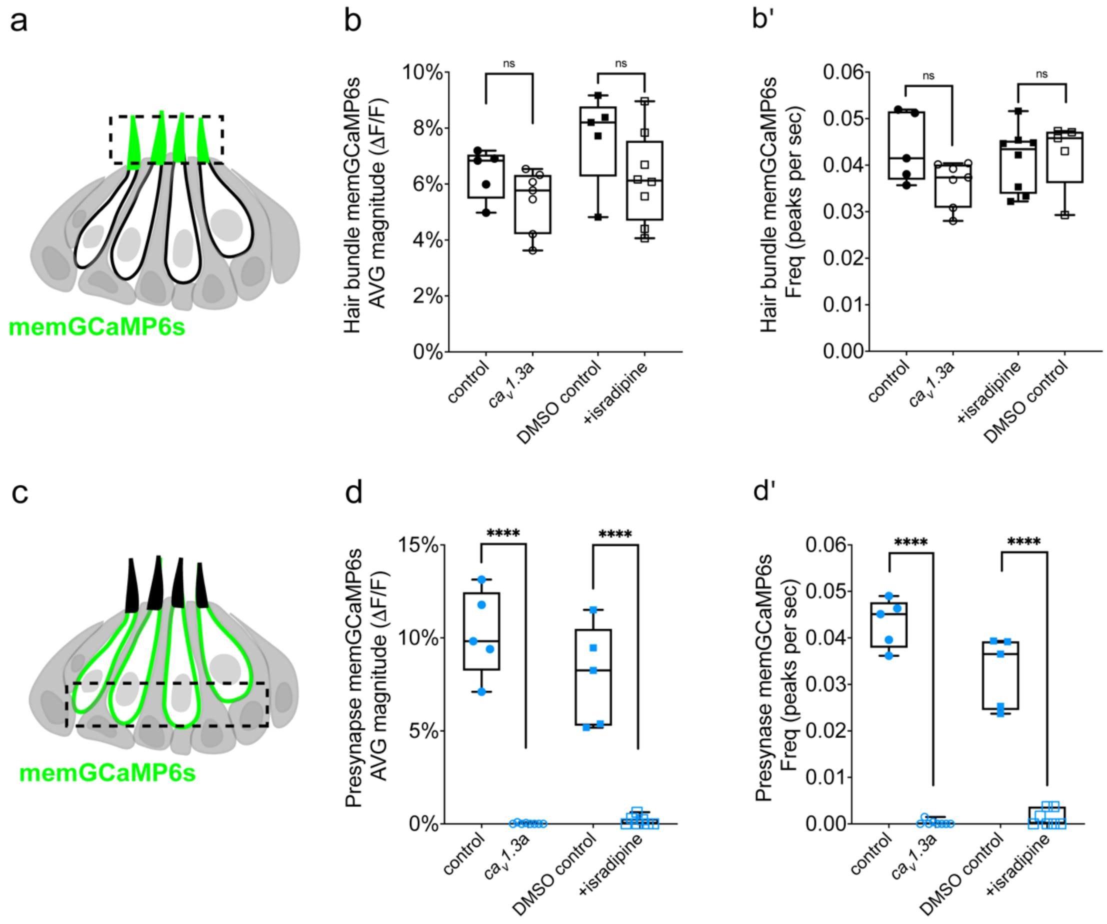


**Figure 5—figure supplement 1. Quantification the change of spontaneous activities in both the hair bundle and presynaptic region after disruption of $Ca_V$1.3a channels.** (a) Diagram illustrating the hair cells with the hair bundles highlighted by a dashed box where we measure their spontaneous calcium activity. (b-b') Box plots show that the average magnitude (b) and frequency (b') of spontaneous calcium activity in the hair bundles is not altered after $Ca_V$1.3a channel disruption using genetic and pharmacological approaches. (c) Diagram illustrating the hair cells with the presynaptic area highlighted by a dashed box where we measure the spontaneous presynaptic calcium activity. (d-d') Box plots show that the average magnitude (d) and frequency (d') of spontaneous calcium activity at the presynaptic regions is completely blocked after $Ca_V$1.3a channel disruption using genetic and pharmacological approaches. All measurements were performed in immature hair cells at day 3. Each point in b-b' and d-d' represents one neuromast. An unpaired t-test was used in b-b' and d-d'. ****p<0.0001.

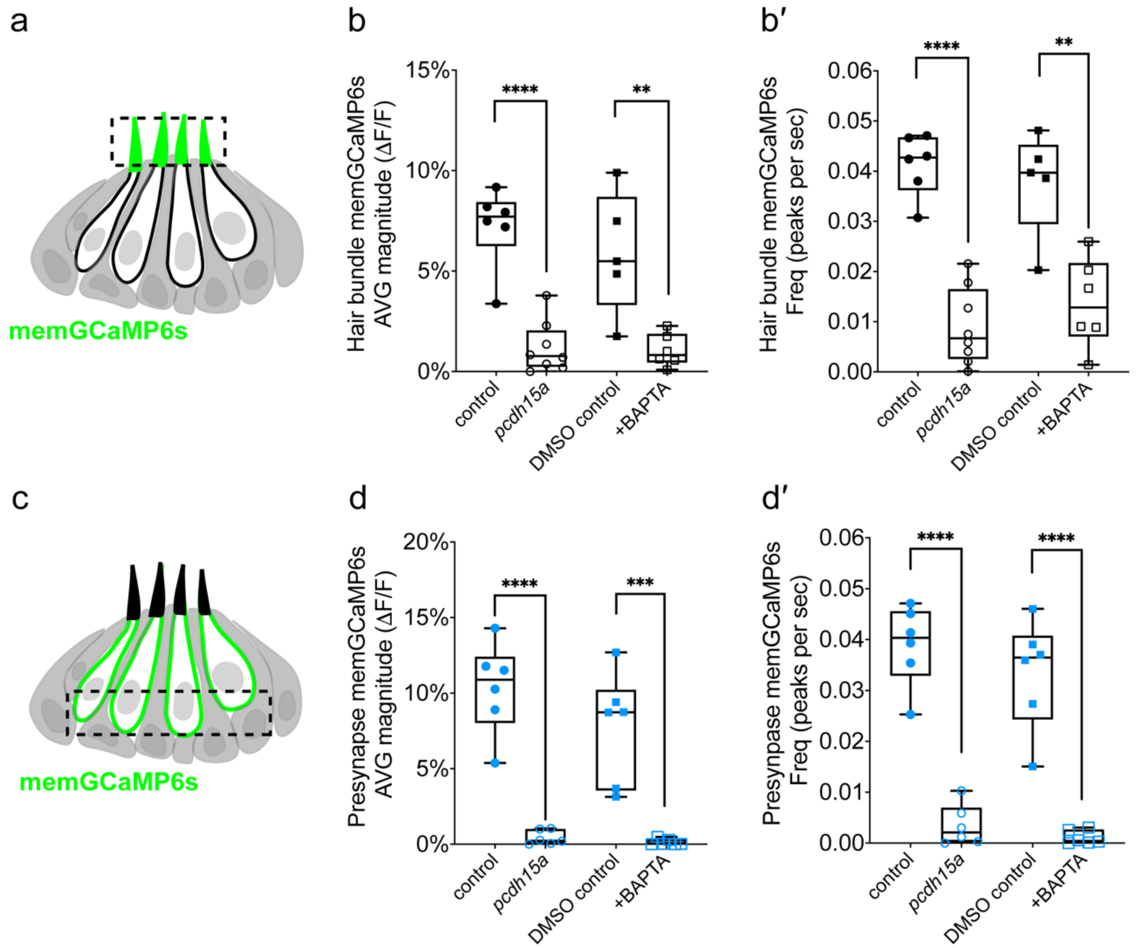


**Figure 6—figure supplement 1. Quantification the change of spontaneous activity in both the hair bundle and presynapse region after mechanotransduction disruption.**

(a) Diagram illustrating the hair cells with the hair bundles highlighted by a dashed box where we measure their spontaneous calcium activity. (b-b') Box plots show the average magnitude (b) and frequency (b') of spontaneous calcium activity in the hair bundles with and without disruption of mechanosensitive function using genetic and pharmacological approaches. (c) Diagram illustrating the hair cells with the presynaptic area highlighted by a dashed box where we measure the spontaneous presynaptic calcium activity. (d-d') Box plots show the average magnitude (d) and frequency (d') of spontaneous calcium activity at the presynaptic regions with and without disruption of mechanosensitive function using genetic and pharmacological approaches. All measurements were performed in immature hair cells at day 3. Each point in b-b' and c-d' represents one neuromast. An unpaired t-test was used in b-b' and d-d'. **p = 0.01, ***p = 0.001, ****p<0.0001.

### Movie Legends

**Movie 1: Volumetric calcium spontaneous activity in developing hair cells**

Volumetric calcium spontaneous activity in the developing hair cells in the lateral line neuromast of a living zebrafish detected using cytoGCaMP6s at day 3. Increases in GCaMP6s intensity are indicated by the heatmap where increases are indicated in red. Images were acquired at 3 s/volume for 900 s using our diSPIM microscope. Playback is 90× real time.

**Movie 2: Spontaneous calcium activity in developing hair cells**

Spontaneous calcium activity in a developing lateral line neuromast of a living zebrafish detected using cytoGCaMP6s at day 3. Images were acquired at 3 s/volume for 900 s using our diSPIM microscope. A single plane is shown. The grayscale images have been converted to the FIJI lookup table 16 colors. Increases in GCaMP6s intensity are indicated by yellow, red and white. Playback is 90× real time.

**Movie 3: Spontaneous calcium activity in supporting cells**

Spontaneous calcium activity in a developing lateral-line neuromast of a living zebrafish using cytoGCaMP6s at day 3. Images were acquired at 3 s/volume for 900 s using our diSPIM microscope. A single plane is shown. The grayscale images have been converted to the FIJI lookup table 16 colors. Increases in GCaMP6s intensity are indicated by yellow, red and white. Playback is 90× real time.

**Movie 4: Two-color spontaneous calcium activities in hair cells and supporting cells**

Two-color spontaneous calcium activities in all hair cells (cytoRGECO1, magenta) and supporting cells (cytoGCaMP6s, green) within a whole neuromast organ at day 3 (Same neuromast as shown in ***Figure 1 and Figure 1–Supplemental Figure 2***). The two signals were acquired simultaneously using our diSPIM microscope at 5 s/volume for 900 s. A single plane is shown. Playback is 150× real time.

**Movie 5: 2-color spontaneous calcium activities in hair cells and contacting efferent terminals**

Two-color spontaneous calcium activities in all hair cells (cytoRGECO1, magenta) and efferent terminals (cytoGCaMP6s, green) within a whole neuromast organ at day 3 (Same neuromast as shown in ***Figure 3***). The 2 signals were acquired simultaneously using our diSPIM microscope at 5 s/volume for 900 s. A single plane is shown. Playback is 150× real time.

**Movie 6: Spontaneous calcium activity in mechanosensory bundles and presynapse within the same hair cells: side by side**

Spontaneous calcium activity in apical mechanosensory bundles (left side) and presynaptic area (right side) of the same hair cells are shown side by side (Same neuromast as shown in ***Figure 4***). MemGCaMP6s was used to detect these calcium signals in lateral-line hair cells at day 2. Images were acquired at 3 s/volume for 900 s using our diSPIM microscope. 2 planes are shown, side by side. The grayscale images have been converted to the FIJI lookup table 16 colors. Increases in GCaMP6s intensity are indicated by yellow, red and white. Playback is 90× real time.

**Supplementary Table 1**

| | SPIM A | | SPIM B | | After Deconvolution | |
|---|---|---|---|---|---|---|
| | Lateral FWHM (µm) | Axial FWHM (µm) | Lateral FWHM (µm) | Axial FWHM (µm) | Lateral FWHM (µm) | Axial FWHM (µm) |
| 1 | 0.44 | 1.36 | 0.435 | 1.445 | 0.33 | 0.395 |
| 2 | 0.52 | 1.55 | 0.44 | 1.42 | 0.24 | 0.415 |
| 3 | 0.41 | 1.415 | 0.435 | 1.89 | 0.31 | 0.48 |
| 4 | 0.46 | 1.595 | 0.435 | 1.525 | 0.38 | 0.425 |
| 5 | 0.44 | 1.385 | 0.48 | 1.385 | 0.345 | 0.47 |
| 6 | 0.495 | 1.445 | 0.46 | 1.545 | 0.35 | 0.32 |
| 7 | 0.55 | 1.465 | 0.44 | 1.49 | 0.305 | 0.4 |
| 8 | 0.42 | 1.565 | 0.47 | 1.54 | 0.295 | 0.375 |
| mean | **0.47** | **1.47** | **0.45** | **1.53** | **0.32** | **0.41** |
| std | 0.05 | 0.08 | 0.02 | 0.15 | 0.04 | 0.05 |
